# *Medicago truncatula CORYNE* modulates inflorescence meristem branching, nutrient signaling, and arbuscular mycorrhizal symbiosis

**DOI:** 10.1101/2024.09.20.614181

**Authors:** Johnathan Orosz, Erica Xinlei Lin, Yerisf C. Torres Ascurra, Mikayla Kappes, Penelope Lindsay, Sagar Bashyal, Hasani Everett, Chandan Kumar Gautam, David Jackson, Lena Maria Müller

## Abstract

The CLAVATA signaling pathway regulates plant development and plant-environment interactions. CLAVATA signaling consists of mobile, cell-type or environment-specific CLAVATA3/ESR-related (CLE) peptides, which are perceived by a receptor complex consisting of leucine-rich repeat receptor-like kinases such as CLAVATA1 and receptor-like proteins such as CLAVATA2, which often functions with the pseudokinase CORYNE (CRN). CLAVATA signaling has been extensively studied in various plant species for its role in meristem maintenance and in legumes for modulating root interactions with nitrogen-fixing rhizobia. Some signaling proteins involved in development and nodulation, including CLAVATA1, also regulate plant interactions with mutualistic arbuscular mycorrhizal (AM) fungi. However, our knowledge on AM symbiosis regulation by CLAVATA signaling remains limited and only a handful of genetic regulators have been identified. Here we report that *Medicago truncatula CRN* controls inflorescence meristem branching and negatively regulates root interactions with AM fungi. *MtCRN* functions partially independently of the AM autoregulation signal *MtCLE53*. Transcriptomic data revealed that *crn* roots display signs of perturbed nutrient, symbiosis, and stress signaling, suggesting that *MtCRN* plays various roles in plant development and interactions with the environment.

## Introduction

Cell-cell communication is critical for plant development and cellular responses to internal and external cues. CLAVATA (CLV) signaling, which consists of mobile CLAVATA3/ESR-related (CLE) peptide hormones and CLE receptor complexes, regulates many aspects of plant development and responses to the environment (Bashyal *et al*., 2023; Lindsay *et al*., 2023). CLE receptor complexes consist of leucine-rich repeat receptor-like kinases (LRR-RLKs) with an extracellular LRR ligand-binding domain and an intracellular kinase domain (e.g., CLAVATA1, CLV1, or BARELY ANY MERISTEM, BAM1-3), receptor-like proteins (e.g., CLAVATA2, CLV2) containing an extracellular ligand-binding domain but lack a kinase domain, and other components such as the pseudokinase CORYNE (CRN) (Hazak and Hardtke, 2016). In Arabidopsis and maize, CLV2 and CRN are considered a CLE perception unit (Bleckmann *et al*., 2009; Je *et al*., 2018*a*). CLV2-CRN can form complexes with CLV1 or BAMs, but also function independently as a parallel pathway (Guo *et al*., 2010; Somssich *et al*., 2016b; Hazak *et al*., 2017). Similarly, the *Medicago truncatula* ortholog of CLV1 (termed SUNN in this species), MtCLV2 and MtCRN can interact with each other (Crook *et al*., 2016). While the cellular function of CRN is still elusive, co-expression assays in *Nicotiana benthamiana* leaves suggest that the short extracellular domain of AtCRN is important for stabilizing the CRN-CLV2 complex at the plasma membrane, whereas the intracellular kinase domain is important for transmitting CLE peptide signals (Somssich *et al*., 2016a).

Mutations in CLV receptors and ligands disrupt meristem organization, often leading to enlarged vegetative, inflorescence and floral shoot meristems (Lindsay *et al*., 2023). These meristem defects in turn impact floral organization and fruit development, altering the number of flowers produced in an inflorescence, the number of floral organs produced, or the arrangement and size of the fruit. The impact of a particular CLV mutation on meristem organization depends on the specific gene mutated, the nature of the mutation (allele), and the plant species; this phenomenon is thought to be a result of differences in compensatory mechanisms in meristem maintenance among plant species (Rodriguez-Leal *et al*., 2019). To date, maize and Arabidopsis are the only plant species in which *crn* mutants have been described. In both species, shoot and inflorescence meristems are enlarged. Arabidopsis *crn* mutants produce more carpels per flower, and some flowers make stamens without anthers (Müller *et al*., 2008). Depending on the temperature, *crn* floral primordia and flowers are prematurely terminated (Jones *et al*., 2021). Maize *crn* mutants produce fasciated ears, with a disordered arrangement of kernel rows and larger kernel row number (Je *et al*., 2018a). CLV signaling also plays roles in root development, including maintenance of the root apical meristem and protophloem differentiation (Stahl *et al*., 2013; Araya *et al*., 2014; Hazak *et al*., 2017). *AtCRN* is expressed in multiple cell types in the root, including stele, endodermis, and quiescent center cells of the meristem (Somssich *et al*., 2016a; Hazak *et al*., 2017). Phloem-expressed AtCRN was shown to be required for the perception of root-active CLE peptides (Hazak *et al*., 2017). Despite its importance for CLE signaling not all Arabidopsis *crn* mutant alleles show a phenotype characterized by longer primary roots (Müller *et al*., 2008; Somssich *et al*., 2016a; Hazak *et al*., 2017), suggesting CRN by itself is not required for root development.

In addition to plant development, CLAVATA signaling pathways also modulate plant interactions with the environment, which includes responses to abiotic and biotic factors such as host responses to pathogenic or beneficial microbes (Bashyal *et al*., 2023). Mutualistic plant interactions with rhizobia during nodulation or *Glomeromycotina* fungi during arbuscular mycorrhiza (AM) symbiosis are regulated by various receptor-ligand signaling pathways (Roy and Müller, 2022), including CLV signaling. In both symbioses, the host plant provides microbial symbionts with photosynthetically fixed carbon in exchange for mineral nutrients. While rhizobia supply the host with nitrogen (N), AM fungi provide a range of benefits including N, phosphorus (P), and water (Smith and Reid, 2008). Both symbioses are governed by negative regulatory pathways based on plant nutrient (N or P) status and prior microbial colonization (autoregulation); these pathways are mediated by nutrient- or symbiosis-induced CLE peptides, respectively (Roy and Müller, 2022). Autoregulation of nodulation (AON) is a systemic negative feedback loop that has been studied in numerous legume species (Wang *et al*., 2018). In *M. truncatula*, AON is regulated by root-derived, nodule-induced CLE peptides (MtCLE12, MtCLE13), which negatively regulate nodule number in a SUNN- and MtCRN-dependent manner (Schnabel *et al*., 2005; Crook *et al*., 2016; Nowak *et al*., 2019). Grafting experiments revealed that SUNN and MtCRN act in the shoot (Schnabel *et al*., 2005; Crook *et al*., 2016); however, both genes are also expressed in roots (Schnabel *et al*., 2012; Crook *et al*., 2016), suggesting various functions across tissues and organs. Autoregulation of mycorrhizal symbiosis (AOM) is regulated by the CLE peptide MtCLE53, which is induced in the vascular tissue of colonized roots and negatively regulates AM fungal colonization by repressing biosynthesis of strigolactones (Müller *et al*., 2019; Karlo *et al*., 2020). Strigolactones are produced by host plants in response to P and N starvation and promote AM symbiosis (Yoneyama *et al*., 2007; Gomez-Roldan *et al*., 2008; Liu *et al*., 2011; Foo *et al*., 2013; Marro *et al*., 2022). In the current AOM model, AM-induced *MtCLE53* down-regulates strigolactone biosynthesis and consequently reduces subsequent AM fungal root colonization (Müller *et al*., 2019).

Like AON, AOM depends on CLV1, which has been shown to regulate AM fungal root colonization not only in legumes (e.g., *M. truncatula* SUNN), but also in non-legumes including tomato and the monocot *Brachypodium distachyon*, suggesting conservation of the pathway across plant clades (Wang *et al*., 2018, 2021; Müller *et al*., 2019; Wulf *et al*., 2024). Plants defective in *SUNN* display a hypermycorrhizal phenotype with elevated root colonization levels and arbuscule numbers and are insensitive to *MtCLE53* overexpression (Morandi *et al*., 2000; Müller *et al*., 2019; Karlo *et al*., 2020). Although AON and AOM signal through the same LRR-RLK SUNN, the upstream-acting CLE peptides and downstream signaling appear to at least partially differ between the two symbioses (Mortier *et al*., 2010; Müller *et al*., 2019; Karlo *et al*., 2020; Chaulagain *et al*., 2023b,a). Recent evidence from tomato (*Solanum lycopersicum*) suggests that AM symbiosis is regulated by multiple, at least partially independent CLV signaling pathways: grafting experiments revealed that while SlFAB (ortholog of CLV1) acts in the root, SlCLV2 is part of two distinct signaling pathways controlling AM symbiosis from the shoot and the root, respectively (Wang *et al*., 2021). In addition, the AM-induced SlCLE11 peptide acts as a negative regulator of AM symbiosis but functions independently of SlFAB or SlCLV2 (Wulf *et al*., 2024). By contrast, hypernodulating pea mutants in *CLV2* (*sym28*), did not show a hypermycorrhizal phenotype, suggesting species-specific differences in symbiosis regulation (Morandi *et al*., 2000). In addition, in contrast to *MtCLE53* over-expression, *sunn* itself (as well as mutants in the tomato and pea orthologs) does not exhibit mis-regulated strigolactone biosynthesis (Foo *et al*., 2014; Müller *et al*., 2019; Karlo *et al*., 2020; Wang *et al*., 2021), suggesting additional signaling components must be involved in MtCLE53-SUNN-mediated AOM signaling that impact strigolactone biosynthesis.

In addition to autoregulation, AM symbiosis is also tightly regulated by the availability of soil mineral nutrients to the plant. High N or P concentrations in the substrate can suppress AM symbiosis development (Abbott *et al*., 1984). Although P is the predominant element plants acquire through AM fungi, N starvation can promote AM symbiosis even under high P (Nouri *et al*., 2014). In *M. truncatula*, P-induced *MtCLE33* acts as a negative regulator of AM symbiosis via a SUNN-dependent pathway although SUNN itself is not required for P repression of AM symbiosis (Müller *et al*., 2019). Similarly, work in tomato revealed that regulation of AM by CLV receptors is independent of plant P but dependent on plant N status (Wang *et al*., 2021). Despite substantial progress in recent years, our knowledge on AM symbiosis regulation by CLV signaling is incomplete and fragmented across plant species. Like with meristem development, CLV signaling may be differently wired across plant species with different nutrient uptake strategies and capacities for symbiotic interactions.

Here, we report *MtCRN* as a regulator of *M. truncatula* inflorescence branching. In addition, we find that *MtCRN* acts as a negative regulator of AM symbiosis in *M. truncatula*. Our results further suggest that, while *MtCRN* contributes to AOM, it also regulates AM symbiosis via additional pathways. Transcriptomics data suggest that *crn* mutants display broad alterations in response to biotic and abiotic factors, indicating that MtCRN may integrate multiple signaling pathways with distinct outcomes.

## Results

### *M. truncatula* CLAVATA signaling regulates inflorescence meristem branching and flowering time but is dispensable for root development

Mutants in CLV signaling have previously been described to be impaired in meristem maintenance, with mutants often displaying inflorescence or stem fasciation (Clark *et al*., 1997). Arabidopsis *CRN* specifically has been reported to modulate inflorescence meristem termination and flower outgrowth under stress (Müller *et al*., 2008; Jones *et al*., 2021). *CLV1* orthologs in legumes (*M. truncatula, Lotus japonicus*, pea, soybean), which regulate symbiosis, show no shoot apical meristem defects (Searle *et al*., 2003; Schnabel *et al*., 2005; Mirzaei *et al*., 2017; Scott *et al*., 2024). Intriguingly, soybean is the only one of these species that has a second CLV1 gene (*GmCLV1A*); *GmCLV1A* does not regulate symbiosis but modulates shoot branching as well as flower number and development, suggesting functional diversification of the two paralogs (Mirzaei *et al*., 2017). In addition, other symbiosis-regulating CLV signaling components such as *LjCLV2, PsCLV2*, and *LjKLAVIER*, as well as the signaling peptide LjCLV3, affect the size of shoot and inflorescence meristems (Oka-Kira *et al*., 2005; Miyazawa *et al*., 2010; Krusell *et al*., 2011; Okamoto *et al*., 2011; Scott *et al*., 2024). Together, this suggests that in legumes, CLV-mediated symbiosis and developmental signaling pathways appear to operate at least partially independently.

To further dissect CLV signaling during legume development, we investigated above-ground developmental phenotypes of *M. truncatula* mutants carrying a homozygous Tnt1 insertion within the coding sequence of the pseudokinase MtCRN (Crook *et al*., 2016). Time to flowering appeared to be delayed in *crn* when compared to R108 wildtypes (Supplementary Fig.1). Stem fasciation, which is characteristic for many CLV pathway mutants in other species (Somssich *et al*., 2016b), was not observed. *M. truncatula* typically produces one solitary flower per inflorescence unit (Fig. 1A, B) (Benlloch *et al*., 2003). However, we noted that *crn* mutants displayed two flowers per peduncle with high frequency: 91% of 8-week old *crn* (n= 11 inflorescences from 3 independent plants) produced two flowers (often with an additional third bud appearing from the same inflorescence, Fig. 1C, D, Supplementary Fig. 1), whereas R108 wildtype plants of the same age and grown under the same growth conditions only produced a single flower per peduncle (n=14 inflorescences from 3 independent plants, Fig. 1A, B, Supplementary Fig. 1). Notably, similar inflorescence phenotypes (76% of inflorescences, n=25) were detected in *M. truncatula* mutants defective in the *CLV1* ortholog *SUNN* (*sunn*-5; Supplementary Fig. 1), suggesting both *SUNN* and *MtCRN* modulate inflorescence meristem branching. Although *crn* mutants produced two flowers per peduncle, ultimately only a single seed was produced per inflorescence as one of the two flowers typically appeared to terminate prematurely. In Arabidopsis, *crn* mutants show increased numbers of floral organs and increased inflorescence meristem size (Müller *et al*., 2008). However, manual dissection of flowers as well as scanning electron microscopy analysis of inflorescences revealed that floral organ numbers and inflorescence width of *M. truncatula crn* did not differ from those in R108 wildtypes (Fig. 1 A-G). This indicates that, unlike

### Arabidopsis and maize, *M. truncatula crn* alters meristem branching, resulting in increased flower number, rather than stem cell proliferation

In roots, CLV signaling is critical for root meristem maintenance, lateral root emergence, and protophloem differentiation (Hazak and Hardtke, 2016). We set out to investigate if *M. truncatula crn* mutants play a role in root development by analyzing root system architecture of 12-day old *crn* and R108 seedlings grown on agar plates. *M. truncatula crn* mutants did not display strong defects in root system development in our experimental growth conditions. We did not detect differences in primary root length, lateral root number, lateral root length, or lateral root density (lateral roots per cm primary root length) between *crn* and R108 seedlings (Supplementary Fig. 2). Similarly, no differences in root length and fresh weight between 4-week-old *crn* and R108 were observed (Supplementary Fig. 2). This is reminiscent of Arabidopsis, where *crn* mutants do not show altered root development although CRN is required for the perception of root-active CLE peptides (Müller *et al*., 2008; Hazak *et al*., 2017).

**Figure 1:**
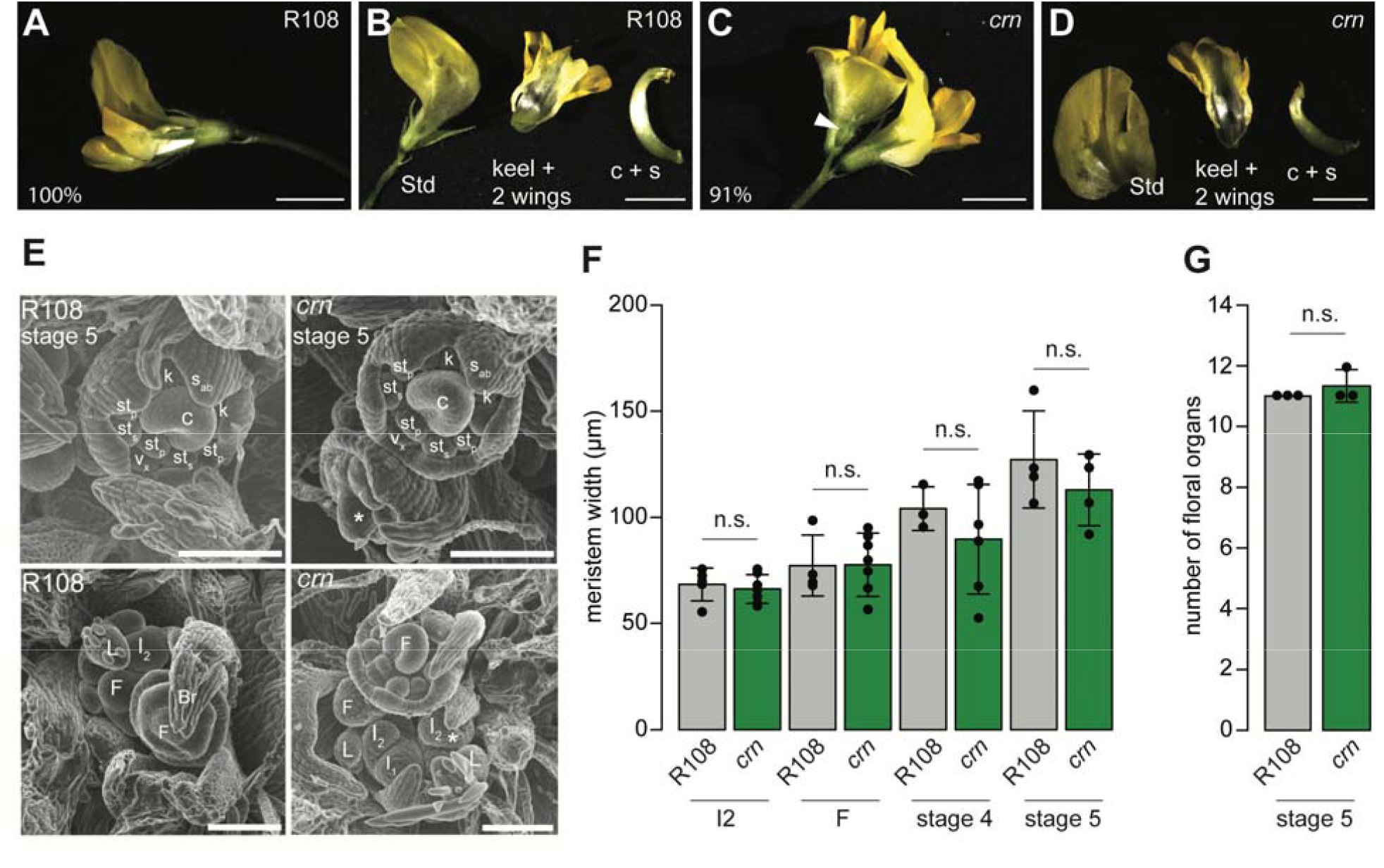
*M. truncatula* CRN impacts inflorescence development. A) Close-up of R108 wild-type flower. 100% of inflorescences produced a solitary flower in our growth conditions. B) Dissected wild-type flower. Std, standard petal; fused keel and 2 wing petals; c+s, carpel and stamen. C) Close-up of inflorescence produced by *crn*. Note the additional undeveloped bud (arrowhead). In our growth conditions, 91% of *crn* mutants produced two flowers per inflorescence, the remaining 9% produced solitary flowers similar to the one shown in (A). D) Dissected *crn* flower. Flower organ number and morphology appeared similar to the R108 wildtype shown in (B). (A-D) Scale bars: 0.5 mm. E) Scanning electron microscopy images of R108 and *crn* inflorescence apices. In *M. truncatula*, the primary inflorescence meristem (I_1_) produces a secondary inflorescence meristem (I_2_) and leaf primordium (L). The secondary meristem will in turn produce a floral meristem (F), which will differentiate floral organ primordia. Top panels show stage five floral meristems, in which all floral organ primordia have been established. Floral organ primordia size and number are unaltered in *crn*, but there is a second stage 5 floral meristem, indicated with an asterisk. The bottom panel shows a range of floral meristem stages in R108 and *crn*. In *crn*, two I_2_ meristems are present at the same time, indicated by an asterisk. C, carpel primordium, k, keel petal, s_ab_, abaxial sepal, st_p_, non-fused vexillary stamen, st_s_, antesepal stamen, v_x_, vexillum, Br, bract. Scale bar: 100 µm. F) Width of R108 and *crn* meristems in different floral meristem stages. No effect of the *crn* mutation was observed on meristem width in any developmental stage assessed. I_2_, secondary inflorescence meristem, F, floral meristem, in which sepal primordia begin to form, stage 4, in which common primordia that will form stamens and petals are formed, stage 5, in which all floral organ primordia have been established. G) Number of stamen and petal floral organ primordia in stage 5 meristems. N.s., no significant difference between R108 and *crn* (Student’s t-test).

### MtCRN negatively regulates AM symbiosis

*M. truncatula* CRN was previously described as a negative regulator of nodulation symbiosis in concert with SUNN (Crook *et al*., 2016; Nowak *et al*., 2019). AON, AOM, and nutrient regulation of AM symbiosis share multiple common components, including SUNN, but the role of CRN in AM symbiosis regulation has not been investigated. Because tomato clv2 mutants are hypermycorrhizal (Wang *et al*., 2018), CRN-CLV2 are commonly considered a signaling unit (Müller *et al*., 2008), and MtCRN can interact with the AOM regulator SUNN as well as MtCLV2 (Crook *et al*., 2016), we hypothesized that *MtCRN* may also regulate root interactions with AM fungi. Indeed, *crn* mutants displayed significantly increased overall Rhizophagus irregularis root colonization levels relative to the R108 wildtype (Fig. 2A). A detailed analysis revealed that *crn* mutants also show higher arbuscule density (i.e., number of arbuscules per mm colonized root in a defined region at the hyphopodium; see methods) than wildtype roots, whereas vesicle density (vesicles/mm colonized roots) did not differ between the wild-type and the mutant (Fig. 2B-E). Intriguingly, similar phenotypes were observed in *Zea mays* (maize) *crn* mutants (Je *et al*., 2018b), which showed a slight increase in overall root colonization and significantly higher arbuscule density than B73 wildtype controls (Supplementary Fig. 3). Together, these results are consistent with the hypothesis that CRN acts as a negative regulator of AM symbiosis and that this role is conserved across plant clades.

**Figure 2:**
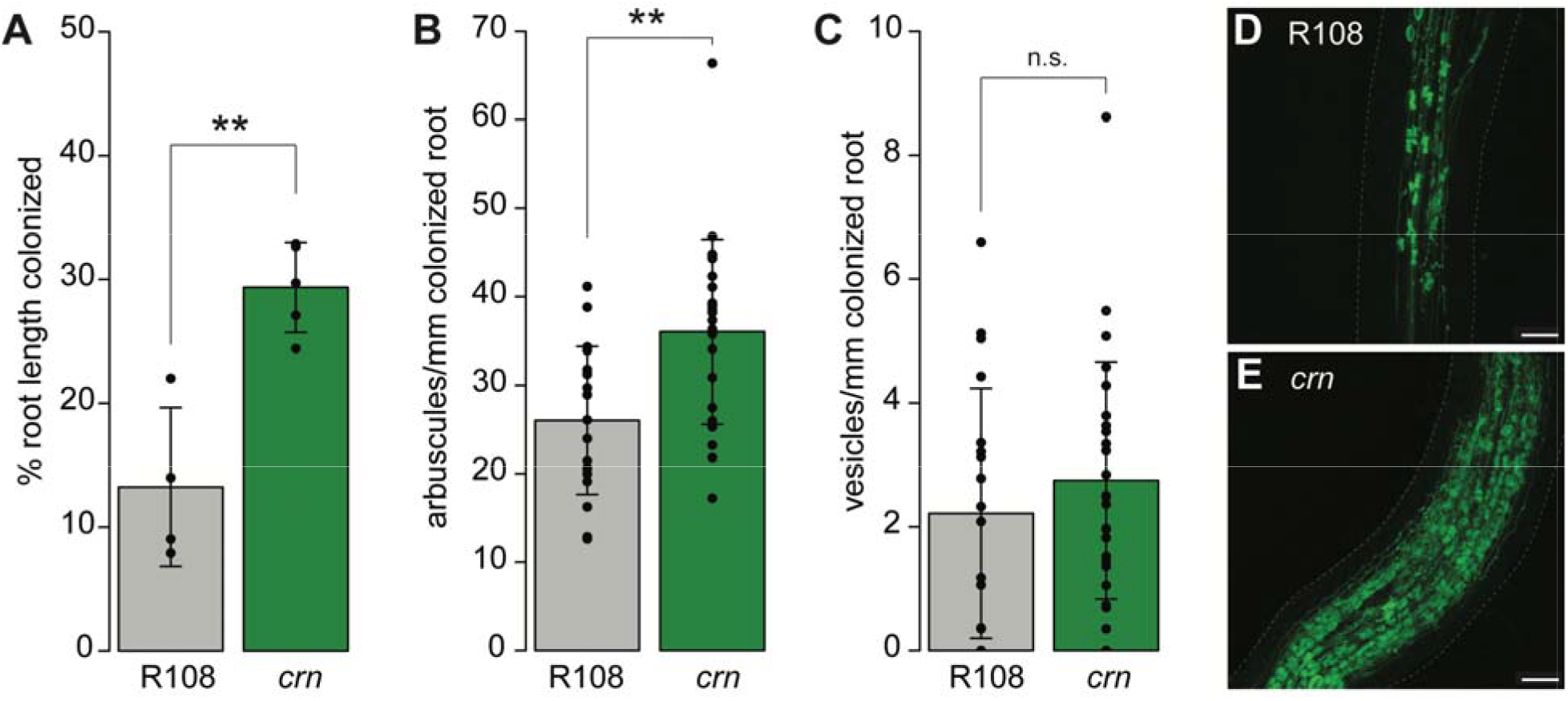
*M. truncatula crn* mutants display elevated levels of colonization by the AM fungus *R. irregularis*. A) *crn* mutants display higher levels of overall root length colonization relative to R108 wild-type controls. B) *crn* mutants show higher arbuscule density (number of arbuscules per mm colonized root length) relative to R108 wild-type controls. C) Vesicle density (vesicles per mm colonized root length) does not differ between R108 and *crn*. D) Representative image of an *R. irregularis*-colonized R108 wildtype root. *R. irregularis* was stained with WGA-Alexafluor488 (green). Dashed line: outline of the root. E) Representative image of an *R. irregularis* (green) colonized root section in *crn*. Dashed line: outline of the root. A-C) Bar plots show averages, error bars the standard deviation, points represent individual measurements (i.e., n=4 individual root systems per genotype in A) and n=19 (R108) and n=25 (*crn*) imaged root sections in B) and C), derived from the same root systems as shown in A). Graphs display data from one experiment; this experiment was repeated twice with similar results. ** p<0.01 (t-test). Scale bar in D), E): 100 µm.

### *MtCRN* is expressed in colonized and non-colonized roots

To gain further insight into the mechanisms of MtCRN regulation of AM symbiosis, we tested the spatial expression pattern of the gene in roots. To do this, we cloned a promoter-reporter construct (*pMtCRN::GUS*) using ~1.9 bp sequence upstream of the *MtCRN* transcription start site and transformed it into A17 wild-type roots using hairy root transformation (Boisson-Dernier *et al*., 2001). In mock-inoculated roots, we observed strong GUS staining in the vasculature and root tips, including lateral root primordia, as previously observed (Crook *et al*., 2016) (Supplementary Fig. 3). In roots colonized by the AM fungus *R. irregularis*, GUS activity was also observed in colonized cortex cells and adjacent, non-colonized cortex cells, in addition to expression in the vasculature and root tip (Fig. 3A, Supplementary Fig. 3). Although our experiment only allowed us to test spatial expression patterns in roots, transcriptomic data suggest that *MtCRN* is also highly expressed in above-ground tissues (Karlo *et al*., 2020; Cope *et al*., 2022). Together, this expression pattern suggests that *MtCRN* plays roles in multiple cell types and tissues. In the context of AM symbiosis, the vascular and cortical promoter activity suggests that *MtCRN* may be involved in the perception of local and systemic CLE signals.

**Figure 3:**
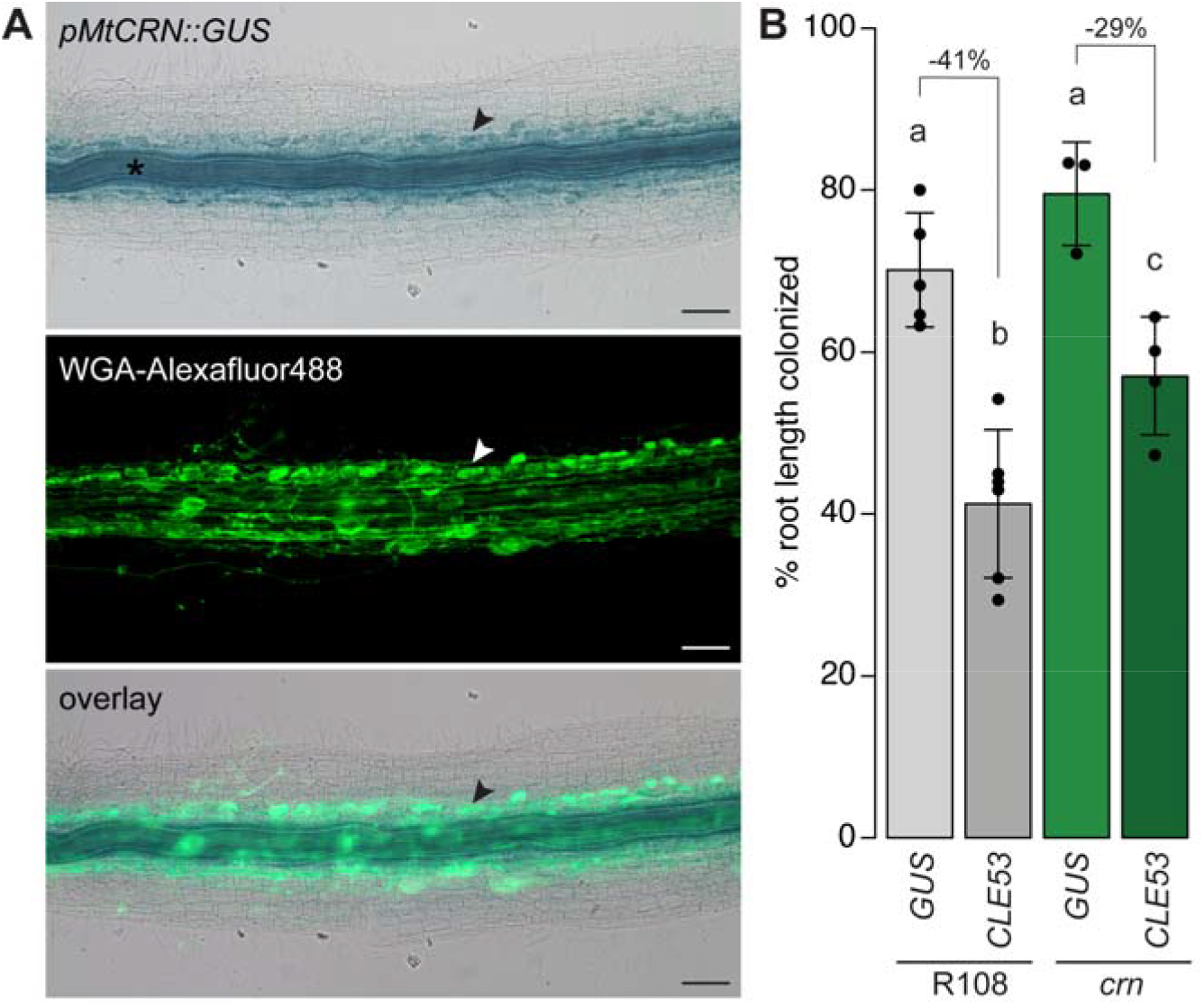
Expression pattern of *MtCRN* in roots and connection to AOM pathway. A) Spatial expression pattern of *pMtCRN::GUS* in *R. irregularis*-colonized roots (top); WGA-Alexafluor488 counterstain to visualize *R. irregularis* (middle); overlay of GUS and WGA-Alexafluor488 images (bottom). GUS activity is observed in the vascular tissue (asterisk) and in colonized cortex cells containing arbuscules (arrowhead). Scale bars: 100 µm. B) AM fungal root length colonization of *crn* mutants partially depend on *MtCLE53*. Graph depicts average root length colonization (%) of R108 and *crn* plants transformed with the *35S::GUS (GUS)* or the *35S::MtCLE53 (CLE53)* construct (compare Supplementary Fig. 3C). *MtCLE53* over-expression reduces *R. irregularis* root colonization in R108 and *crn*; however, this suppressive effect is stronger in R108 than in *crn*. For each genotype, the % of colonization reduction upon MtCLE53 over-expression is shown. The bar graph shows average values; error bars denote standard deviation. Black dots indicate individual datapoints (biological replicates; n=3-5 replicates per sample). This graph displays data from one experiment; the experiment was repeated twice with similar results. Different letters denote significant differences (p<0.05) after ANOVA (p=2.12*10^−5^) followed by Tukey’s HSD test for pairwise comparisons.

### *MtCRN* functions partially independent of the AOM signal MtCLE53

We previously reported that *MtCRN* is required for *MtCLE16* and *RiCLE1* signaling, plant and fungal CLE genes induced in cortex cells upon colonization by AM fungi, respectively (Bashyal *et al*., 2025). The *pMtCRN::GUS* signal detected in colonized cortex regions suggests that cortical MtCRN may be involved in a local response to these CLE peptides. However, our GUS data also suggest high *MtCRN* expression in the vascular tissue, which is indicative of additional roles in other, vascular or systemic CLE signaling pathways. Autoregulation of nodulation (AON) and mycorrhizal symbiosis (AOM) negatively regulates nodulation and AM symbiosis, respectively, based on existing symbioses (Wang *et al*., 2018). We previously reported that ectopic over-expression of the *M. truncatula* AOM signal MtCLE53, which is expressed in the vascular tissue, reduced overall root length colonization in a SUNN-dependent manner (Müller *et al*., 2019). MtCRN has also been implicated in the systemically-acting AON and functions downstream of the nodulation-induced CLE peptides MtCLE12 and MtCLE13 (Nowak *et al*., 2019). To test if *MtCRN* also acts down-stream of AOM signals, we over-expressed *MtCLE53* under the control of the constitutively active Cauliflower mosaic virus 35S promoter (*35S::MtCLE53*) as well as a control construct (*35S::GUS*) in roots of *crn* mutants and R108 wild-type controls (Supplementary Fig. 3). Similar to what was previously reported for the *M. truncatula* A17 accession (Müller *et al*., 2019; Karlo *et al*., 2020), ectopic over-expression of *MtCLE53* in roots also reduced overall AM fungal root length colonization in R108. Interestingly, unlike *sunn*, which is fully insensitive to *MtCLE53* over-expression suggesting the LRR-RLK acts in the same pathway as the peptide hormone (Müller *et al*., 2019), ectopic *MtCLE53* overexpression reduced AM fungal root length colonization also in *crn* (Fig. 3B). However, colonization reduction in *35S::MtCLE53* relative to 35S::GUS-expressing roots is weaker in *crn* (29%) than in the wild-type background (41%). The partial sensitivity of *crn* to *MtCLE53* over-expression suggests that *MtCRN* is involved in *MtCLE53* signaling but that the AM symbiosis phenotype of *crn* is also influenced by additional, *MtCLE53*-independent pathways.

### Transcriptomic analysis of *M. truncatula crn* mutants reveals global changes in stress response, nutrient homeostasis, and environment interactions

To understand the molecular mechanisms causing the elevated AM fungal root colonization levels in *crn* mutants, we sequenced the transcriptomes of 3 *R. irregularis*-colonized and 4 mock-inoculated *crn* and R108 wild-type root systems, respectively (Supplementary table 1). As expected, we observed major transcriptomic changes induced by *R. irregularis* colonization in R108 wild-type roots (434 up-regulated and 266 down-regulated genes, here defined as differentially expressed genes with a log2 fold-change >1 or <−1, respectively, and padj<0.05; Supplementary figures 4-6). Comparing *R. irregularis*-colonized *crn* mutant roots with mock-inoculated *crn* roots, we identified 816 up- and 325-downregulated genes (Supplementary figures 4-5,7). In mock-inoculated *crn* roots relative to mock-inoculated wild-type roots, we identified 1506 up- and 647 down-regulated genes (Supplementary figures 4-5, 8). When comparing differentially expressed genes in *R. irregularis*-colonized *crn* roots with *R. irregularis*-colonized wild-type roots, we identified 1061 up- and 594-downregulated genes (Supplementary figures 4-5, 9, Supplementary table 2). Because *crn* roots are more colonized than wild-type roots, these observed transcriptomic differences may be caused by differences in genotype, differences in AM fungal colonization levels, or both. We performed hierarchical clustering analysis of all differentially expressed genes in *R. irregularis*-inoculated *crn* roots relative to *R. irregularis*-inoculated wild-type roots to identify clusters of differentially expressed genes with similar expression patterns and to distinguish differentially expressed genes solely influenced by colonization levels from those regulated by *MtCRN*.

Differentially expressed genes were grouped into six clusters (Fig. 4A, Supplementary table 2). Clusters 1, 3 and 6 contained genes that are regulated by AM fungal root colonization, and whose general expression pattern changes were similar in *crn* and wild-type roots. However, the degree of up- or down-regulation differs between the genotypes, likely caused by the differential colonization levels in the mutant compared to the wildtype (Fig. 2 A-E).

**Figure 4:**
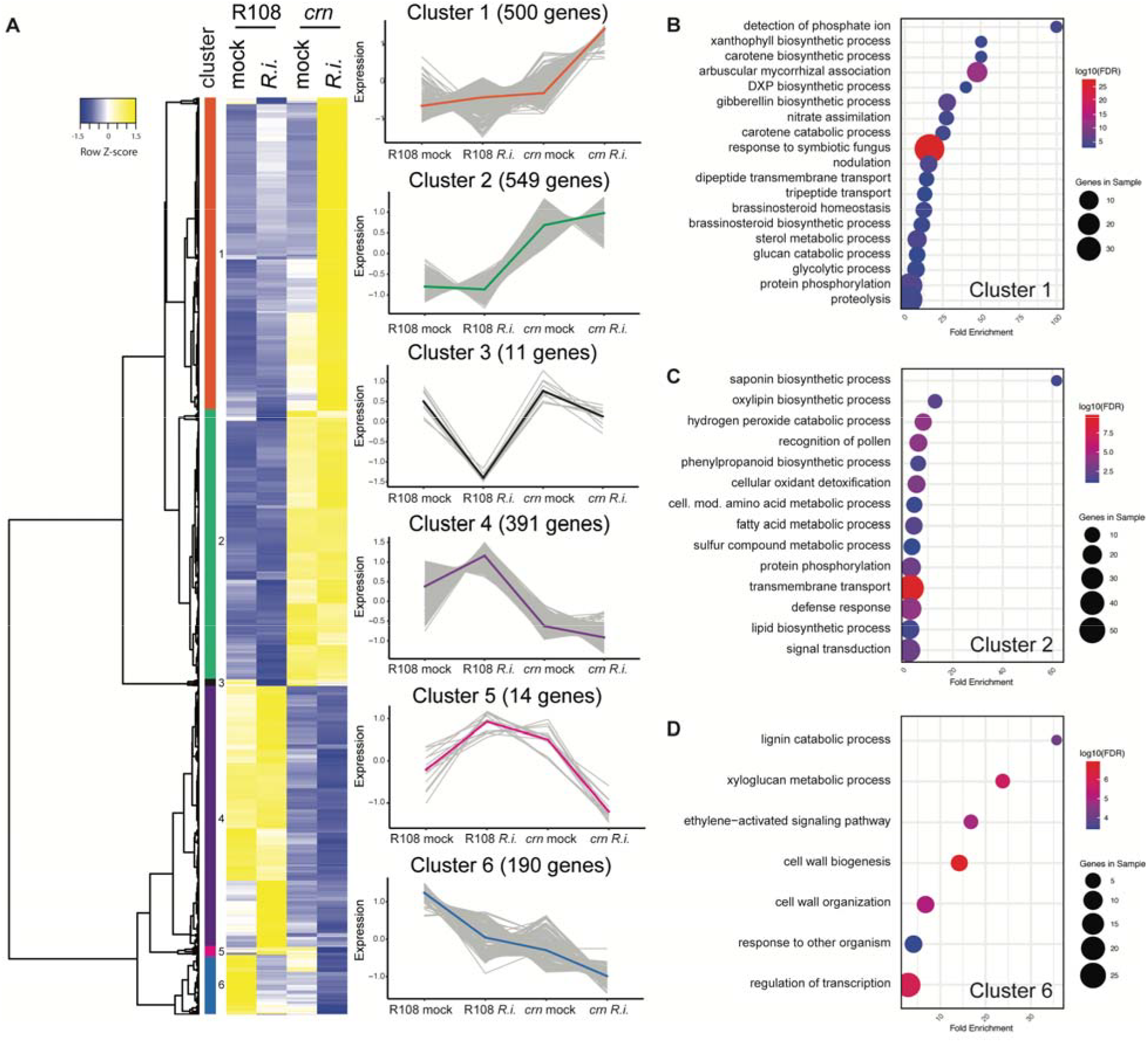
Hierarchical cluster analysis of differentially expressed genes in *R. irregularis*-colonized and mock-inoculated *crn* and R108 wild-type roots. A) 1655 genes that are differentially expressed between *R. irregularis*-colonized *crn* and R108 wild-type roots (log2 fold-change >1 or <−1, padj<0.05) were selected for cluster analysis to identify gene groups with similar expression patterns. Genes were clustered using Pearson correlation. Heatmap shows averages of n=4 root samples each for *crn* and R108 mock-inoculated, and n=3 samples each for *crn* and R108 *R. irregularis*-inoculated (*R*.*i*.). Gene lists for each cluster can be found in table S2. Line graphs display normalized relative expression changes across samples for each individual gene in a cluster (grey) and average changes in color. B) Biological functions GO terms enriched in cluster 1, C) Biological functions GO terms enriched in cluster 2, D) Biological functions GO terms enriched in cluster 6. Note: Cluster 4 has one significantly enriched GO term “plant-type secondary cell wall biogenesis”, which is not shown in this figure. Supplementary tables 2 and 3 contain a list of genes for each cluster and the associated GO terms, respectively. DXP, 1-deoxy-d-xylulose 5-phosphate. FDR, false discovery rate.

Cluster 1 genes are induced by AM symbiosis and are generally more highly expressed in *R. irregularis*-colonized *crn* roots relative to colonized wild-type roots (Fig. 4A). GO analysis of the genes in cluster 1 revealed, among others, an enrichment of genes associated with GO terms related to symbiosis (“AM association”, “response to symbiotic fungus”, “nodulation”), nitrate assimilation, as well as genes associated with the biosynthesis and transport of secondary metabolites and hormones known to regulate AM symbiosis (e.g., carotenoids, gibberellic acid, brassinosteroids; Fig. 4B, Supplementary table 3). This cluster contains well-studied AM marker genes including but not limited to *MtPT4, MtRAM1, MtRAM2*, or *MtSTR*, which function in bidirectional nutrient exchange between the symbiotic partner and whose expression is known to be correlated with AM colonization levels (Serrano *et al*., 2024).

Clusters 3 and 6 contain genes that are down-regulated by AM fungal root colonization; AM-mediated down-regulation is more pronounced in *crn* roots than in wildtypes. While no significant GO terms were found in cluster 3 (11 genes), in cluster 6 we detected enrichment of GO terms related to cell wall biogenesis and remodeling (Fig. 4D, Supplementary table 3) as well as ethylene signaling. Notably, this GO term contained five transcription factors of the APETALA2/ETHYLENE RESPONSE FACTOR (AP2/ERF) family; however, AP2/ERF transcription factors have a more role in plant stress signaling, and, apart from ethylene, also often respond to the hormones abscisic acid (ABA), gibberellic acid (GA), brassinosteroids (BR), and cytokinin (Xie *et al*., 2019). To our knowledge, no symbiosis-related function has been reported for any of the AP2/ERF transcription factors in cluster 6.

Cluster 2 contains 549 genes that are up-regulated in *crn* roots relative to wild-type roots, regardless of AMF colonization status (Fig. 4A). Many genes in this cluster appear to play roles in plant responses to abiotic and biotic stresses: The GO term with the highest fold-enrichment (61-fold) in this cluster is saponin biosynthetic process (Fig. 4C, Supplementary table 3). Saponins are secondary metabolites with antioxidant activity that play various roles in plant-microbe interactions, including plant defense against pathogens and herbivores (Mugford and Osbourn, 2012; Chen *et al*., 2014). Soy saponins have been identified in root exudates as modulators of plant interactions with rhizosphere microbes (Tsuno *et al*., 2018; Fujimatsu *et al*., 2020). In addition, 12 peroxidases associated with the GO terms “hydrogen peroxide catabolic process” and “cellular oxidant detoxification” were enriched in cluster 2. Peroxidases are major regulators of cellular reactive oxygen species (ROS) homeostasis (Passardi *et al*., 2005). ROS play various roles in plants, including response to pathogens and beneficial microbes, including AM, but also development, signaling, and cell growth (Wojtaszek, 1997; Fester and Hause, 2005; Huang *et al*., 2019). The GO term “cellular modified amino acid metabolic process” was enriched in cluster 2; this GO term contained (among others) 4 genes annotated as glutathione S-transferases, which also have antioxidant activity and are involved in ROS scavenging and cellular detoxification during stress responses (Gullner *et al*., 2018). Together, this data suggest that cellular oxidative stress may be attenuated in *crn* through increased ROS scavenging. Conversely, *crn* roots display an enrichment of GO terms associated with lipid metabolism (“lipid biosynthetic process”, “fatty acid metabolic process”, “monocarboxylic acid biosynthetic process”) and oxylipin biosynthesis. The latter contained four genes encoding 9-lipoxigenases (*9-LOX*), and one gene annotated as *ALPHA-DIOXYGENASE 1* (*αDOX1*). 9-oxylipins play a role in cellular signaling, defense, and abiotic stress response (Vicente *et al*., 2012), and their up-regulation suggests that *crn* mutants may have a heightened response to biotic stress even in the absence of microbes. In line with this hypothesis, the GO term “defense response” was also enriched among the genes in cluster 2 (Fig. 4C); this GO term contained several genes encoding NUCLEOTIDE-BINDING SITE LEUCINE-RICH REPEAT (NBS-LRR) family proteins, which can directly bind to microbial effectors and thus play critical roles in plant immune responses to pathogens (Marone *et al*., 2013). This suggests that, despite signatures of attenuated oxidative stress, other components of the plant response system to biotic stresses are up-regulated in *crn*.

Furthermore, we detected an enrichment of genes associated with the GO term “transmembrane transport” among the *crn*-upregulated genes in cluster 2. Interestingly, these included, among others, genes encoding transporters for carbon products such as sugar (*MtESL16*.*1/MtESL2*.*4, MtESL3*.*4, MtESL6*.*4, MtSTP3*.*2, MtSTP5*.*1/MtSTP5*.*2*), which may indicate that carbon flux is altered in *crn*. In addition, we identified putative transporters for nitrate (*MtNPF2*.*11, MtNPF5*.*13, MtNPF5*.*5, MtNPF5*.*10*), ammonium (*MtrunA17_Chr4g0007621/ MtrunA17_Chr4g0007611*), phosphate (*MtPT2, MtPT9*), metals (*MtMTP8*), potassium (MtrunA17_Chr4g0055471), and zinc (MtZIP6) among the genes in cluster 2 (Supplementary table 2, Supplementary table 3), suggesting mis-regulation of various nutrient uptake or homeostasis pathways in the mutant (further discussed below).

Cluster 4 contains genes that are down-regulated in *crn* roots relative to wild-type roots with little impact of symbiosis status (Fig. 4A). In this group of 391 genes, only the GO term ‘plant-type secondary cell wall biogenesis’ was enriched (Supplementary table 3). Intriguingly, all 13 genes associated with this GO term in cluster 4 are annotated as FASCICLIN-LIKE ARABINOGALACTAN PROTEIN (FLA). FLAs are extracellular glycoproteins involved in a plethora of functions, including development, growth, and environmental responses (He *et al*., 2019), but to our knowledge have not been studied in the context of AM symbiosis or CLAVATA signaling. However, genes encoding arabinogalactan proteins as well as other cell wall-related genes have been shown to be down-regulated also in a A. thaliana *crn* mutant (Pallakies and Simon, 2014). This suggests a conserved role of CRN in cell wall organization, which is critical for its function in growth and development.

Cluster 5 contains 14 genes that are induced in R108 *R. irregularis*-colonized roots relative to mock-inoculated roots but show the opposite pattern in *crn* background (down-regulated in *R. irregularis*-colonized roots relative to mock-inoculated *crn* roots) (Fig. 4A, Supplementary table 3). No significantly enriched GO terms were detected in cluster 5.

### N and P homeostasis signaling is perturbed in *crn* roots

Our cluster analysis suggests an up-regulation of multiple mineral nutrient transporters in *crn*, regardless of root colonization status (see above, Supplementary table 3). This suggests that in *crn* mutants, nutrient starvation or homeostasis signaling pathways may be mis-regulated. Because P and N are major regulators of AM symbiosis and their homeostasis is regulated by CLAVATA signaling pathways in various plant species (Bashyal *et al*., 2023), we examined genes associated with P and N homeostasis more closely in our dataset. Interestingly, 16 genes putatively associated with N homeostasis were regulated by *MtCRN* regardless of root colonization status, and 11 additional ones were found to be synergistically regulated by *MtCRN* and AM status of the plant (Fig. 5A-B, Supplementary table 4). Similarly, eight genes associated with P homeostasis were mis-regulated in *crn* relative to R108 without additional regulation by root colonization status, and an additional three genes were regulated by MtCRN and AM symbiosis (Fig. 5C-D, Supplementary table 4). This data suggest a major perturbance in nutrient homeostasis signaling in *crn* even in the absence of symbiosis. Because nutrient starving plants initiate more interactions with beneficial microbes (Nouri *et al*., 2014), a mis-regulation of nutrient homeostasis or starvation signaling may promote AM fungal root colonization in *crn*. To test this hypothesis, we grew *R. irregularis*-inoculated R108 and *crn* plants under varying phosphate and nitrogen conditions (Supplementary Fig. 3). Under low nitrogen (0.5 mM) with low or high phosphate (20µM or 2 mM, respectively), the *crn* mutant phenotype was suppressed and no difference in root length colonization was detected between the mutant and the wildtype. Conversely, when grown under a high nitrogen (50 mM) fertilization and low phosphate fertilization regime, *crn* mutants displayed higher root length colonization than wildtype controls. The sensitivity of the *crn* mutant phenotype to nitrogen levels is reminiscent to what has been reported for mutants in the CLV1 and CLV2 orthologs in tomato (Wang *et al*., 2021), and indicates that MtCRN may participate in a CLAVATA signaling pathway involved in nitrogen regulation of AM symbiosis.

**Figure 5:**
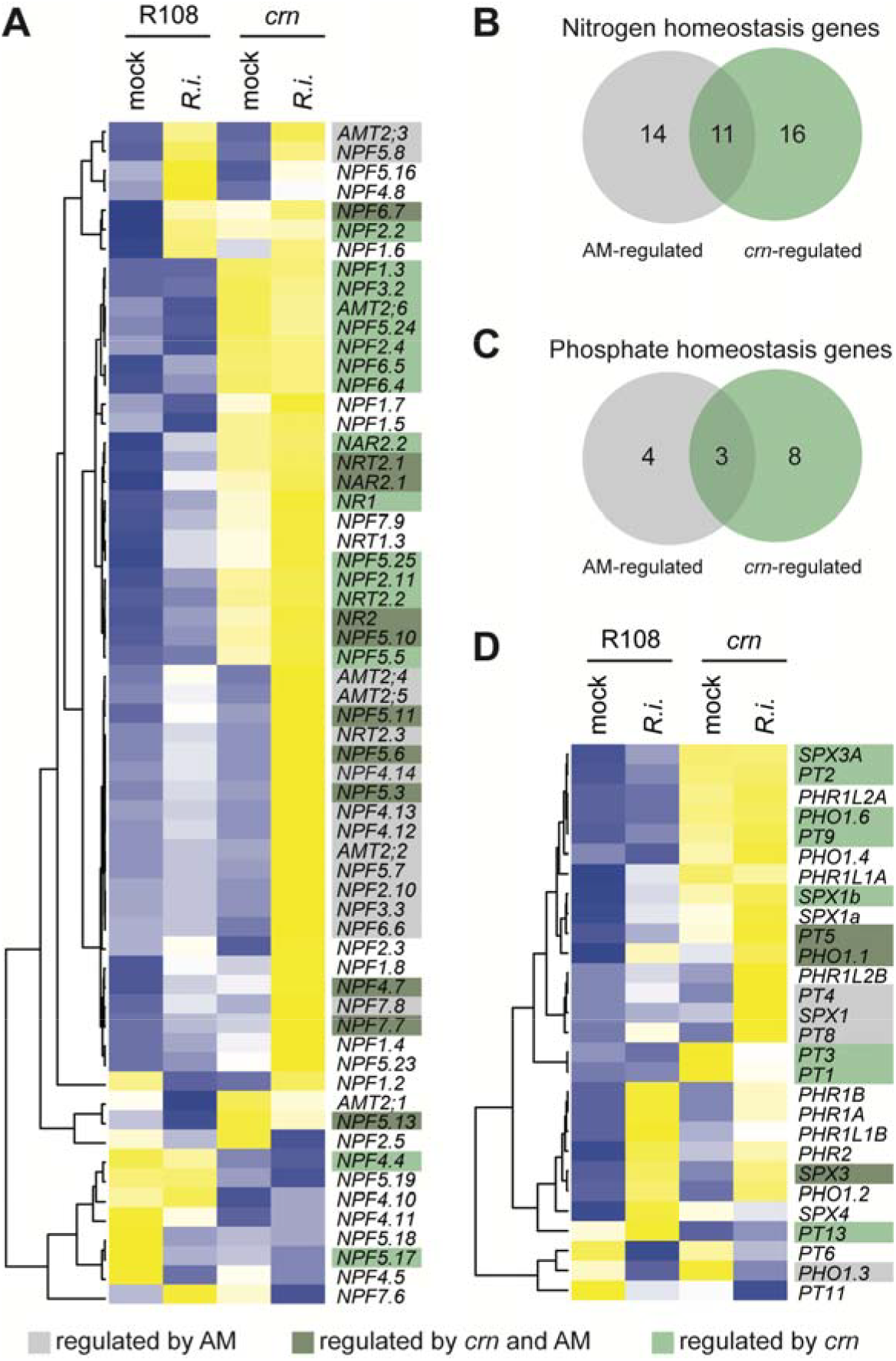
Differential regulation of N and P homeostasis genes in *crn* mutants. A) Heatmap for genes annotated as *NITRATE TRANSPORTER (NRT), NITRATE AND PEPTIDE TRANSPORTER FAMILY (NPF), NITRATE ASSIMILATION RELATED (NAR), AMMONIUM TRANSPORTER (AMT)*, and *NITRATE REDUCTASE (NR)* with putative roles in *M. truncatula* N metabolism. B) Venn diagram showing the number of putative N homeostasis genes responding to AM symbiosis, *crn*, or both (padj<0.05). C) Venn diagram showing the number of putative P homeostasis genes responding to AM symbiosis, *crn*, or both (padj<0.05). D) Heatmap for genes annotated as *PHOSPHATE TRANSPORTER (PT), PHOSPHATE1 (PHO), PHOSPHATE STARVATION RESPONSE (PHR and PHR-LIKE, PHRL)*, or *SYG1/PHO81/XPR1* family (*SPX*) with putative roles in P homeostasis and transport. A and D) Heatmaps show averages of n=4 root samples each for *crn* and R108 mock-inoculated, and n=3 samples each for *crn* and R108 *R. irregularis*-inoculated (R.i.). AM-regulated genes are highlighted in grey, genes differentially regulated in *crn* vs. R108 with no effect of colonization status in light green. Genes responsive to AM and *crn* are highlighted in dark green. B and C) ‘AM-regulated’ is defined as genes with differential regulation (padj<0.05) between R.irregularis-colonized and mock-inoculated R108 and/or *crn* roots, respectively. ‘*crn*-regulated’ is defined as genes with differential regulation (padj<0.05) in mock-inoculated *crn* relative to mock-inoculated R108 roots. Transcript data used to produce these heatmaps can be found in Supplementary table 4.

### Expression of early symbiosis signaling genes is mis-regulated in *crn*

Consistent with an increase in AM fungal root colonization, we observed elevated gene expression of known AM-induced genes in *R. irregularis*-colonized *crn* roots relative to colonized wild-type controls (see above). This was the case for well-studied genes involved in arbuscule accommodation and symbiotic nutrient exchange (e.g., *MtPT4, MtRAM2, MtSTR*), and arbuscule degeneration (e.g., *MtMYB1, MtCP3, MtTGL*) (Fig. 6A and B, Supplementary table 5). Expression of these genes did not differ when mock-inoculated *crn* roots were compared to mock-inoculated R108 roots.

**Figure 6:**
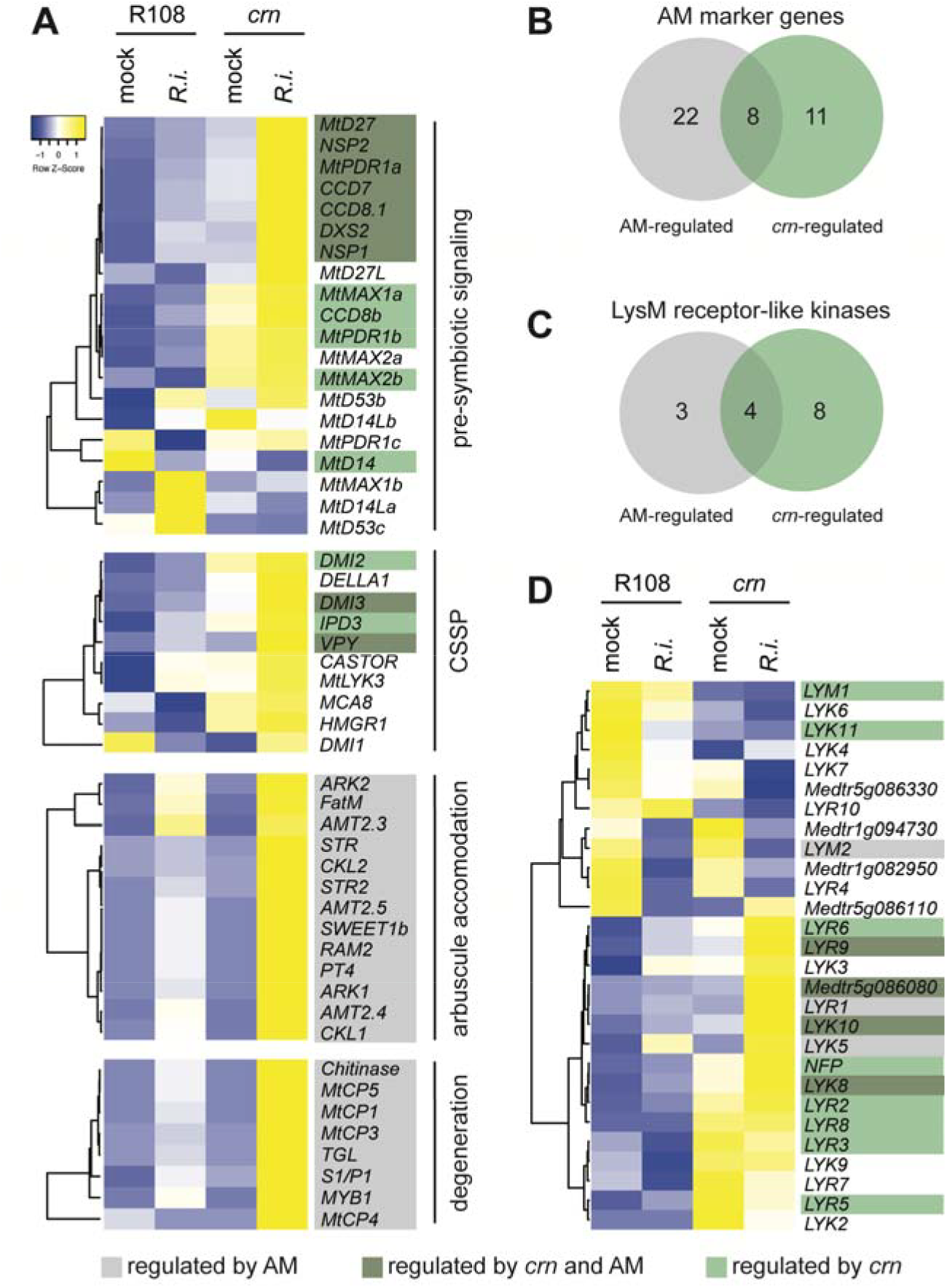
Differential regulation of AM symbiosis regulators in *crn* mutants. A) Heatmaps for known genes involved in pre-symbiotic signaling, the common symbiosis signaling pathway (CSSP) shared between AM and nodulation symbiosis, arbuscule accommodation, and arbuscule degradation. B) Venn diagram showing the number of AM regulatory genes from panel A) responding to AM symbiosis, *crn*, or both. C) Venn diagram showing the number of LysM-RLK or LysM receptor protein-encoding genes (LYK, LYR, LYM) responding to AM symbiosis, *crn*, or both. D) Heatmap for LysM-RLK genes in mock- and *R. irregularis*-inoculated wildtype (R108) and *crn* roots. A and D) Heatmaps show averages of n=4 root samples each for *crn* and R108 mock-inoculated, and n=3 samples each for *crn* and R108 *R. irregularis*-inoculated (R.i.). AM-regulated genes are highlighted in grey, genes differentially regulated in *crn* vs. R108 with no effect of colonization status in light green. Genes responsive to AM and *crn* are highlighted in dark green. B and C) ‘AM-regulated’ is defined as genes with differential regulation (padj<0.05) between *R. irregularis*-colonized and mock-inoculated R108 and/or *crn* roots, respectively. ‘*crn*-regulated’ is defined as genes with differential regulation (padj<0.05) in mock-inoculated *crn* relative to mock-inoculated R108 roots. Transcript data used to produce these heatmaps can be found in supplementary tables 5 and 6.

Conversely, the expression of many genes involved in pre-symbiotic signaling was perturbed in mock-inoculated *crn* roots relative to mock-inoculated wildtype roots, suggesting these genes may be direct or indirect targets of *MtCRN* signaling: We found that the expression of genes involved in strigolactone biosynthesis and secretion (*MtD27, MtCCD7, MtCCD8, MtPDR1a*) not only responds to AM fungal root colonization in *crn* and R108, but is also differentially regulated in *crn* mock-inoculated roots relative to mock-inoculated wild-type controls (Fig. 6A and B). In addition, *MtCCD8b* and *MtMAX1a*, orthologs of genes involved in strigolactone biosynthesis in other species, as well as *MtMAX2* and *MtD14*, involved in strigolactone signaling and perception, respond to AM fungal root colonization and are also differentially regulated in mock-inoculated *crn* roots relative to controls. Similar trends were observed for the known transcriptional regulators governing strigolactone biosynthesis *NSP1* and *NSP2*. Because the production of AM-symbiosis promoting strigolactones is regulated by P and N status of the plant (Yoneyama *et al*., 2007, Yoneyama *et al*., 2012; Liu *et al*., 2011; Foo *et al*., 2013; Marro *et al*., 2022), an up-regulation of the strigolactone biosynthesis pathway may be a consequence of the apparent mis-regulation of N and P homeostasis signaling in *crn* roots (Fig. 5A-D).

In addition, the expression of several genes involved in the common symbiosis signaling pathway (CSSP) appear to be directly or indirectly regulated by MtCRN (Fig. 6A-B). The CSSP is triggered by cellular perception of N-acetylglucosamine (GlcNAc)-derived (lipo)chitooligosaccharide molecules derived from AM fungi or rhizobia (myc-LCOs/COs or nod-LCOs, respectively), and is required for cellular entry of symbiotic microbes at the onset of symbiotic interactions (Oldroyd, 2013). DMI2, which encodes a putative co-receptor perceiving microbial signals, was found to be up-regulated in mock-inoculated *crn* roots when compared to R108. Likewise, *DMI3, IPD3*, and *VPY*, which regulate cellular signal transduction and symbiosis initiation in response to microbial LCO perception (MacLean *et al*., 2017; Lindsay *et al*., 2022) function downstream of MtCRN signaling. In addition, DMI3 and VPY expression was also found to be responsive to AM symbiosis (Fig. 6A). Because the CSSP pathway is not only required for the establishment of AM but also nodulation symbiosis, and because *crn* was reported to have a hypermycorrhizal phenotype (this study) and a hypernodulation phenotype in *M. truncatula* (Crook *et al*., 2016), we hypothesize that *crn* mutants are primed for symbiotic interactions even in the absence of microbes, potentially due to an up-regulation of the symbiosis accommodation program as a result of perturbations in N and P starvation signaling (Li *et al*., 2022). In line with this hypothesis, we found that some genes encoding lysin motif (LysM) RLKs involved in the recognition of nod- and myc-LCOs (e.g., *MtLYK8* and *NFP*, but not *MtLYK3*) are up-regulated in *crn* roots even in the absence of a microbe (Fig. 6 C-D, supplementary table 1, supplementary table 6) (Limpens *et al*., 2003; Smit *et al*., 2007; Zhang *et al*., 2024). Intriguingly, 10 additional LysM RLKs or receptor-like proteins (LYK, LYR, LYM) that can bind LCOs but have no known role in nodulation or symbiosis (Buendia *et al*., 2018) were also up-regulated in *crn* mutants relative to wildtypes (both in *R. irregularis*- or mock-inoculated treatments), suggesting recognition of diverse microbial patterns may be altered in *crn*.

### *Rhizophagus irregularis* transcriptionally responds to colonizing *crn* roots

To investigate *R. irregularis* responses to host genotype, we mapped our transcriptome reads to the *R. irregularis* DAOM197198 v2.0 genome (Yildirir *et al*., 2022). About 1.2-3.8 % of the reads mapped to the fungal genome in colonized R108 and *crn* roots (Supplementary table 1). We found 19 genes to be up-, and 10 genes to be down-regulated in *crn* roots relative to wild-type roots (logFC>1 or <−1, padj<0.05; Fig. 7A, supplementary table 7). Among the functionally annotated genes down-regulated in *R. irregularis* colonizing *crn* roots relative to wild-type roots (logFC<−1, padj<0.05), we identified multiple transcripts putatively involved in cellular signaling, including two protein tyrosine kinases, a C2H2 zinc finger transcription factor, and one SAM-domain containing protein, which mediate protein-protein interactions and have many roles, including cellular signaling (Kim, 2003). This suggests that *R. irregularis* modulates signaling in response to colonizing *crn* roots; however, because AM fungi are functionally not well understood, we cannot predict which signaling pathways may be impacted.

**Figure 7:**
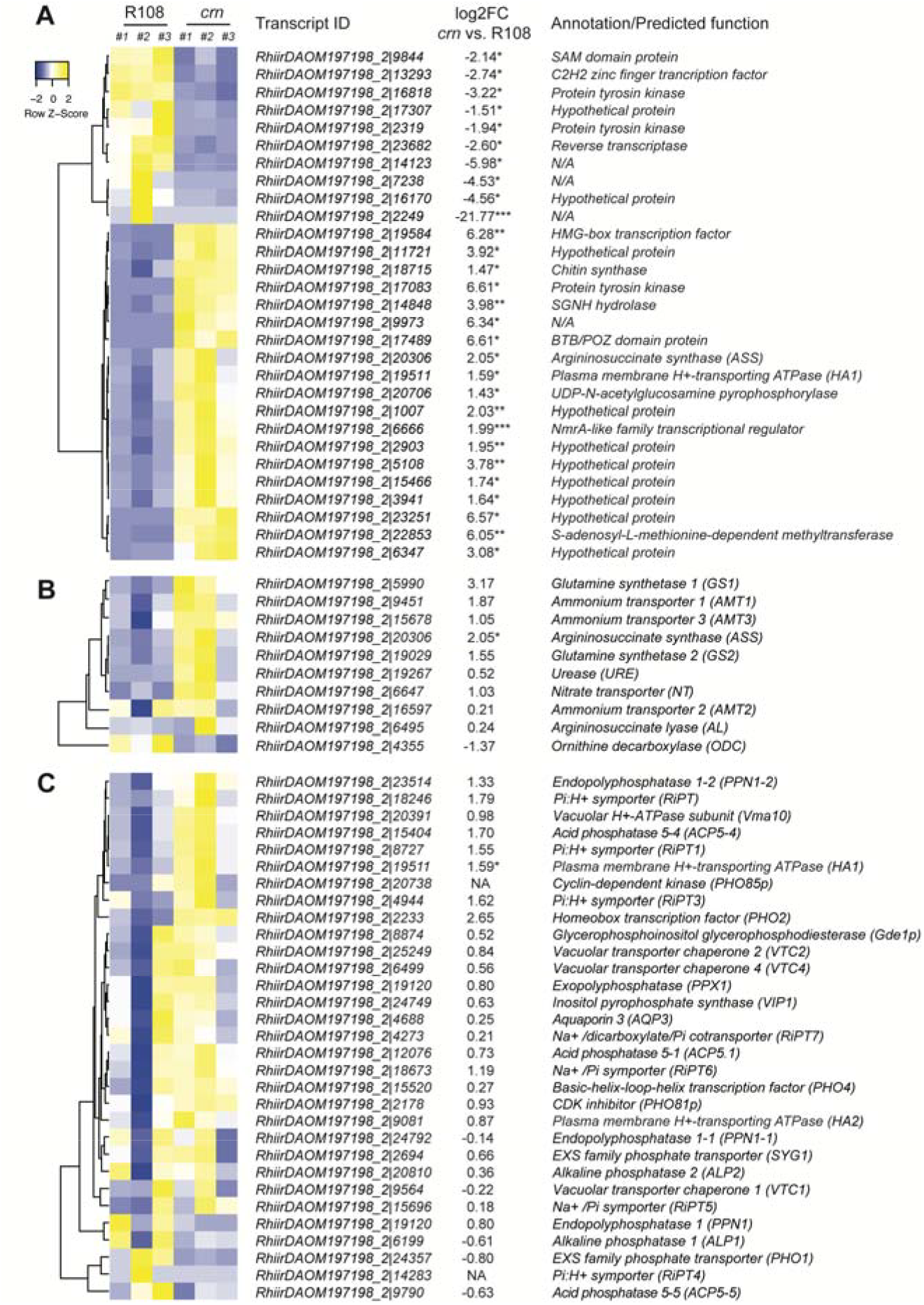
Host genotype-dependent regulation of *R. irregularis* gene expression. A) Differentially expressed *R. irregularis* genes (log2 fold change >1 or <−1, padj<0.05) in *crn* and R108 roots. B) Heatmap of genes involved in fungal N metabolism. C) Heatmap of genes involved in fungal P metabolism. Significance: ***adjusted p-value (padj)<0.05, **padj<0.01, *padj<0.05; if no symbol is shown, fungal gene expression does not significantly differ between host genotypes. Transcript data used to produce these heatmaps can be found in supplementary table 7. Data for the same Argininosuccinate synthase (*ASS*) gene are shown in panels A) and B); data for the same H^+^-transporting ATPase (*HA1*) gene are shown in panels A) and C).

Among the *R. irregularis* genes significantly up-regulated in *crn* roots relative to wild-type roots (logFC>1, padj<0.05), we identified multiple transcripts that suggest fungal N metabolism might be perturbed in *crn* (Fig. 7A). Differentially regulated genes associated with fungal N metabolism include an argininosuccinate synthase (ASS), which is involved in the biosynthesis of arginine, a critical component of N storage in AM fungi (Cruz *et al*., 2007). In addition, among the significantly up-regulated genes we found a transcript annotated as *NmrA (Nitrogen metabolic regulation A)-like protein*, which in other fungi is part of a protein complex that functions in transcriptional regulation of N metabolism based on N availability (Andrianopoulos *et al*., 1998; Stammers, 2001). Both *ASS* and *NmrA* have been predominantly detected in the extraradical mycelium of AM fungi where N is taken up (Dumas-Gaudot *et al*., 2004; Tian *et al*., 2010). Other known genes involved in *R. irregularis* N metabolism tended to be up-regulated in *crn* roots relative to wildtypes (Fig. 7B), although the adjusted p-value did not fall below the stringent significance criteria described above. This included the ammonium transporters GintAMT1 and *GintAMT3*, but not *GintAMT2*, two glutamine synthetases (*GS1, GS2*) and a N transporter (NT) (Fellbaum *et al*., 2012) (Fig. 7B).

Interestingly, a plasma membrane H^+^-transporting ATPase HA1 with suggested roles in fungal P sensing (Xie *et al*., 2022) was also observed among the *R. irregularis* genes significantly up-regulated in *crn* roots (Fig. 7A). This observation prompted us to examine other transcripts known to be involved in fungal in P homeostasis and transport (Kikuchi *et al*., 2014, 2016; Xie *et al*., 2022). Although none of the known P homeostasis or transport genes were differentially regulated in *crn* vs. R108 roots, some genes displayed trends toward increased expression in *crn* relative to R108 (Fig. 7C). This included, among others, the Pi:H^+^ symporter RiPT thought to be involved in fungal P uptake from the environment (Harrison and van Buuren, 1995; Maldonado-Mendoza *et al*., 2001) and the Pi stress marker PHO2 (Austin and Mayer, 2020).

In addition, our *R. irregularis* transcriptome displays signs of increased chitin biosynthesis when colonizing *crn* roots (Fig. 7A, Supplementary table 7). Among the *R. irregularis* genes significantly up-regulated in *crn* relative to R108 roots (logFC>1, padj<0.05), we identified a gene encoding a UDP-N-acetylglucosamine diphosphorylase, which catalyzes the formation of GlcNAc, and a chitin synthase, which catalyzes chitin formation from GlcNAc building blocks (Fig. 7A) (Chen *et al*., 2022). Notably, GlcNAc are also building blocks of LCO and CO molecules critical for pre-symbiotic communication with the host. Myc-LCO and CO production is promoted by host strigolactones (Chaulagain *et al*., 2023a); biosynthesis genes for these phytohormones are induced in *crn* roots relative to wildtypes (Fig. 6), which could be linked to the observed up-regulation of GlcNAc in *R. irregularis*. However, while fungal LCO and CO molecules initiate CSSP signaling in the host upon recognition by plant LysM-type RLKs, they also have a more general function in regulating fungal growth and development (Rush *et al*., 2020). In addition, a member of the SGNH hydrolase/GDSL lipase family was up-regulated. Although the specific function of this gene is unknown, SGNH hydrolases catalyze a variety of carbon-modifying reactions (Anderson *et al*., 2022), which may be related to fungal growth and development within the host plant.

Together, our fungal transcriptome dataset suggests that *crn* roots stimulate *R. irregularis* N and P metabolism, growth and development. Notably, it is possible that some of these changes are an indirect consequence of higher fungal colonization levels observed in the mutant; however, the lack of differential regulation of *R. irregularis* housekeeping genes between *crn* and wild-type roots (Supplementary table 8) suggests that the observed gene expression differences are not solely caused by increased fungal biomass in *crn* roots. While more research is required, our data support the hypothesis that *R. irregularis* differentially responds to as-yet-unknown signals derived from *crn* and wild-type roots.

## Discussion

CLV signaling has been reported to modulate various aspects of plant development and responses to the environment (Bashyal *et al*., 2023; Lindsay *et al*., 2023). Here we describe *M. truncatula* CRN as a central regulator of meristem branching and root interactions with AM fungi.

Although legume mutants in the CLV signaling pathway have been well-studied in multiple plant species in the context of symbiosis regulation, an effect on meristem development has only been described in soybean and pea (Krusell *et al*., 2011; Mirzaei *et al*., 2017; Scott *et al*., 2024). Soybean encodes two CLV1 copies, one of which controls symbiosis and the other controls meristem development (Mirzaei *et al*., 2017). Our work contributes to a better understanding of legume inflorescence meristem patterning by revealing a role for *MtCRN* and *SUNN* in this process. Although the molecular mechanisms remain to be determined, several hints in our data and those of others point toward an intersection of auxin and CLV in *M. truncatula* meristem signaling: Auxin is an important regulator of meristem identity, and AtCRN was found to act upstream of auxin to control floral primordia outgrowth in Arabidopsis (Jones *et al*., 2021). The *M. truncatula* auxin transport mutant *smooth leaf margin1* has pleiotropic developmental effects, including multiple flowers per inflorescence that are reminiscent of the *crn* flower phenotype reported here (Zhou *et al*, 2011), suggesting that MtCRN may control auxin-mediated outgrowth in *M. truncatula* floral development similar to what was found in Arabidopsis (Jones *et al*., 2021). In addition, previous studies reported that *SUNN* regulates shoot-to-root auxin transport to control N response (Jin *et al*., 2012) and that auxin signaling, among other hormonal pathways, is mis-regulated in *sunn* (Karlo *et al*., 2020), further supporting the hypothesis that CLV signaling may act upstream of auxin also in the context of *M. truncatula* meristem maintenance. Like in Arabidopsis, legume meristem signaling may be controlled by multiple, functionally redundant pathways and/or buffering. Consequently, phenotypes may only be apparent in certain mutant alleles or growth conditions. Because these buffering mechanisms appear distinct among plant species, this may explain why inflorescence meristem branching is primarily affected in Medicago *crn* mutants, resembling the phenotype in other legumes (Krusell *et al*., 2011; Mirzaei *et al*., 2017), but inflorescence meristem size is affected in maize and Arabidopsis *crn* mutants (Rodriguez-Leal *et al*., 2019, Hazak and Hardtke, 2016, Je *et al*., 2016).

CLV signaling has been previously described to be critical for autoregulation of AM symbiosis (AOM), but our knowledge on the molecular mechanisms remains limited. We previously characterized the AM-induced, vascular tissue-expressed MtCLE53 peptide as a member of the AOM pathway ((Müller *et al*., 2019; Karlo *et al*., 2020). *MtCLE53* requires *SUNN* to regulate AM symbiosis (Crook *et al*., 2016; Müller *et al*., 2019; Karlo *et al*., 2020). Because MtCRN and SUNN can interact and are required to perceive nodulation-associated CLE peptides during AON (Crook *et al*., 2016; Nowak *et al*., 2019), we hypothesized that MtCRN may also be required for MtCLE53 signaling together with SUNN. However, the data presented here suggest that the negative effect of *MtCLE53* overexpression on AM symbiosis is only partially dependent on *MtCRN*. One explanation for our observation is that MtCLE53 can signal through MtCRN-SUNN but also parallel, MtCRN-independent pathway(s). Such parallel pathways and compensation mechanisms are common in CLAVATA signaling modules (Pallakies and Simon, 2014; Nimchuk, 2017; Je *et al*., 2018a; Rodriguez-Leal *et al*., 2019; Selby and Jones, 2023). Our data also suggests that additional, MtCLE53-independent pathways contribute to the elevated AM fungal root colonization observed in *crn*. One of these additional pathways is likely the one regulated by *MtCLE16*, an AM-induced, cortex-expressed, *M. truncatula* CLE peptide that locally promotes AM symbiosis in concert with MtCRN (Bashyal *et al*., 2025). The *MtCRN* promoter activity, which we detected in colonized cortex cells and the vascular tissue, also supports the hypothesis that the pseudokinase plays a dual role in AM symbiosis control through cortical MtCLE16 and vascular MtCLE53 signaling. Multiple CLV signaling pathways also appear to govern AM symbiosis in tomato (*Solanum lycopersicum*); for example, SlCLV2 was reported to regulate AM symbiosis through two distinct local and systemic signaling pathways (Wang *et al*., 2021; Wulf *et al*., 2024). Notably, the receptor complexes involved in vascular and cortical CLAVATA signaling have not been resolved for AM symbiosis. It is known that CRN can interact not only with CLV1 (SUNN) but also other receptors, including CLV2 (Bleckmann *et al*., 2009; Crook *et al*., 2016; Je *et al*., 2018b) and BAM3 (Hazak *et al*., 2017). Interestingly, MtBAM2 was recently reported to play a role during nodulation symbiosis (Thomas and Frugoli, 2024). Future research should investigate if the same or related RLKs function during AM symbiosis in concert with MtCRN.

Our transcriptomics experiment revealed that *M. truncatula crn* mutants display signatures of abiotic stress, most notably nutrient starvation as suggested by a transcriptional up-regulation of N, P and other nutrient transporters. Intriguingly, *R. irregularis* also displays signs of disrupted N and P metabolism as indicated by an up-regulation of *NmrA* and *ASS*. The proteins encoded by these genes are predicted to function in the extracellular mycelium of AM fungi to mediate N utilization and incorporation into arginine (Andrianopoulos *et al*., 1998; Cruz *et al*., 2007; Fellbaum *et al*., 2012). Arginine plays a key role in N translocation from the extra-to the intraradical mycelium in AM fungi (Cruz *et al*., 2007; Fellbaum *et al*., 2012). In addition, *R. irregularis HA1* expression was increased when colonizing *crn* roots; *RiHA1* was previously implicated in P signaling and uptake by the extraradical mycelium, where it powers Pi:H^+^ symporters to promote phosphate uptake (Ezawa and Saito, 2018; Xie *et al*., 2022). Together with our host transcriptomics data, this finding suggests that *crn* roots may represent a greater sink for N and P than wild-types and consequently stimulate N and P uptake in the colonizing *R. irregularis* symbiont (Cruz *et al*., 2007). Further research is required to better understand the impact of host genotype and host CLV signaling on symbiont metabolism.

In the plant, N and P starvation results in the upregulation of genes involved in the biosynthesis of strigolactones, which in turn act as positive regulators of AM symbiosis (Akiyama *et al*., 2005; Yoneyama *et al*., 2007; Gomez-Roldan *et al*., 2008; Liu *et al*., 2011; Foo *et al*., 2013, 2014; Marro *et al*., 2022). Intriguingly, our data reveal that many of these genes are upregulated in *crn* relative to wildtype controls, which may be a direct consequence of the apparent perturbations in N and P signaling in *crn*. Strigolactone biosynthesis is also the target of AOM, and the elevated expression of strigolactone biosynthesis genes may also be caused by an at least partial disruption of AOM. In either case, upregulation of genes involved in the symbiosis-promoting strigolactones in *crn* can explain the elevated root colonization levels in the mutant. Together, our functional and transcriptomic data suggest that MtCRN may act upstream of strigolactone biosynthesis regulation in response to MtCLE53 as well as N and P signaling. Intriguingly, up-regulation of strigolactone biosynthesis genes have not been observed in CLV1 mutants in *M. truncatula* (sunn), pea (nark) or tomato (fab) (Foo *et al*., 2014; Karlo *et al*., 2020; Wang *et al*., 2021), suggesting that CRN (but not SUNN/FAB) may modulate cross-talk between N and P homeostasis, MtCLE53-mediated autoregulation, and AM fungal root colonization.

Apart from genes involved in nutrient homeostasis and strigolactone biosynthesis, our transcriptome analysis also suggests that *crn* roots display an upregulation of genes associated with the CSSP. The CSSP is required for cellular entry of the microbial symbionts and is activated upon perception of diffusible LCO or CO signals derived from AM fungi or rhizobia (Oldroyd, 2013). Because *crn* has previously been described as a regulator of nodulation symbiosis, an effect of *crn* on CSSP gene expression may contribute not only to the hypermycorrhizal phenotype described here, but also to the hypernodulation phenotype described previously (Crook *et al*., 2016). It was reported previously in rice that transcription of certain CSSP genes (including orthologs of *MtVPY, DMI2, DMI3, IPD3*, which we found here to be regulated by *crn*) is regulated by the plant phosphate starvation response (Das *et al*., 2022). In addition, expression of LysM-type RLKs and LysM receptor proteins as well as plant responses to LCO/CO perception depend on N and P status of the plant and are greatly increased upon nutrient starvation (Li *et al*., 2022). We therefore hypothesize that the perturbed N and P signaling in *crn* promotes AM symbiosis by increasing the expression of genes involved in early symbiosis signaling (strigolactone biosynthesis genes, CSSP genes, LysM-RLKs), which facilitates colonization by AM fungi upon contact. It has been established that nutrient starvation determines the outcome of plant-microbe interactions by fine-tuning plant immune responses, potentially in order to facilitate symbiotic interactions (Paries and Gutjahr, 2023). Consistent with this model, our dataset suggests attenuated oxidative stress in *crn* relative to wildtypes; oxidative stress is a hallmark of responses to abiotic factors but also plant immunity. However, we also detected up-regulation of plant defense genes, suggesting *crn* displays additional signatures of enhanced response to biotic stress. Intriguingly, CLV signaling – including CRN – has been implicated in plant responses to diverse abiotic and biotic stresses, including beneficial and pathogenic microbes (Bashyal *et al*., 2023; Torres Ascurra and Müller, 2025). For example, *A. thaliana crn* mutants display increased resistance to *Heterodera schachtii* cyst nematodes as well as certain bacterial and oomycete pathogens (*Ralstonia solanacearum* and Hyaloperonospora arabidopsidis, respectively), while being more susceptible to Botrytis necrotrophic fungi or *Pseudomonas syringae* bacterial infection (Replogle *et al*., 2011; Hanemian *et al*., 2016). Together with our data, this suggests that coordination or crosstalk of multiple combined abiotic and biotic stress responses, rather than overall stress signaling, may be impaired in *crn*.

In summary, our findings suggest that *MtCRN* plays multiple, likely independent, roles in plant development, nutrient homeostasis, and plant interactions with the environment. Based on data from other plant species, where different combinations of CLV1-type receptors, CLV2, and CRN are responsible for the perception of diverse CLE signals in a context- and cell-type-specific manner (Whitewoods, 2021), we propose that *MtCRN* acts downstream of diverse CLE peptides in the context of meristem development, nutrient homeostasis, and plant-microbe interactions. More research is needed to identify additional context-dependent CLE signals and receptors that provide specificity to the various signaling pathways *MtCRN* is involved in.

## Materials and methods

### Plant materials and growth conditions

Homozygous *M. truncatula crn* (NF3436) and *sunn-5* (NF2262) mutants, both in R108 background, were a gift from Dr. Julia Frugoli (Clemson University, SC, USA) and were previously described (Crook *et al*., 2016; Nowak *et al*., 2019; Thomas and Frugoli, 2024). For AM inoculation experiments, seeds were extracted from seed pods and scarified using sandpaper or by incubating the seeds in concentrated sulfuric acid for 7 minutes. Seeds were washed 5 times in autoclaved distilled H2O and surface-sterilized with 10% Bleach for 7 minutes. Seeds were rinsed again 5 times in autoclaved distilled H2O and incubated at 4°C for 48 hours. Imbibed seeds were germinated on germination paper (Anchor paper) for 3-5 days at room temperature before transplanting into substrate. Plants were grown as previously described (Chaulagain *et al*., 2023a). In brief, germinated seedlings were transplanted into SC10 cone-tainers (Steuwe and Sons) filled with washed and autoclaved substrate mix consisting of play sand and fine vermiculite (1:1 ratio). For inoculated samples, 250 *Rhizophagus irregularis* DAOM197198 spores (Premier Tech, Canada) were placed onto a sand layer 5 cm below the surface of the substrate in each cone-tainer. For mock-inoculated samples, an equal amount of distilled water was added onto the sand layer. Plants were grown in a growth chamber under a 16 hour light (24°C)/8 hour dark (22°C) regime at 40% relative humidity. For standard experiments, all cones were fertilized twice per week with 20 ml half-strength Hoagland’s fertilizer supplied with 20µM potassium phosphate and 2.5 mM nitrate (potassium nitrate and calcium nitrate). For nutrient manipulation experiments, plants were watered twice weekly with 10 ml of half-strength Hoagland’s solution, which was adjusted as follows: nitrate levels were reduced to 0.5 mM and phosphate levels were either 20µM (low nitrogen/low phosphate) or 2 mM (low nitrogen/high phosphate). For high nitrogen/high phosphate, we added 50mM nitrogen and 2mM phosphate, respectively. Concentrations of all other nutrients remained constant. Unless otherwise noted, all plants were harvested 5-6 weeks after inoculation. To enrich for *R. irregularis* colonization, we sampled 5 cm-long root sections centered around the inoculated region. For the transcriptomics experiment, whole root systems were harvested.

### Transient root transformation

Transient root transformations of *M. truncatula* R108 and *crn* plants with *35S:GUS* and *35S:MtCLE53* constructs were performed as described previously (Boisson-Dernier *et al*., 2001; Müller *et al*., 2019). Chimeric plants were inoculated with *R. irregularis* as described above and harvested 6.5 weeks after planting into cone-tainers.

### Root system phenotyping

Seeds were extracted from seed pods and sterilized as described above. Seeds were then distributed on the bottom of a glass petri dish using water surface tension (Floss *et al*., 2013). Glass petri plates were placed upside down in a 28°C incubator and seedlings were allowed to germinate overnight. With this method, we were able to obtain seedlings with straight roots. Seedlings were then transferred to 12×12 cm square petri dishes filled with modified Fahraeus (F) medium supplied with 4 g/L Gelzan CM agar (Sigma-Aldrich) (Floss *et al*., 2013). Plates were placed at a 45° angle in a growth chamber under a 16 hour light (24°C)/8 hour dark (22°C) regime. The bottom 2/3 of the plates was wrapped in aluminum foil to protect the roots from light. Pictures of seedlings were taken every 3 days for 12 days starting with the day of transfer to agar plates (day 0). Root system parameters (primary root length, lateral root length, lateral root number, lateral root density) were analyzed using the SmartRoot package in Fiji (Lobet *et al*., 2011; Schindelin *et al*., 2012) for each timepoint.

### GUS staining

The ~1.9 kb MtCRN promoter sequence was amplified with Phusion High-Fidelity DNA Polymerase (Thermo Scientific) from genomic DNA of wildtype A17, using the primers indicated in Supplementary Table 9. The *pMtCRN:GUS* construct was created by assembling the *MtCRN* promoter, ß-Glucuronidase, GUS, and the destination vector pKm34GW-RR (Floss *et al*., 2017) by Gibson cloning using the NEBuilder HiFi DNA Assembly Master Mix. *M. truncatula* A17 roots were transformed and inoculated with *R. irregularis* as described above and harvested 5 weeks after planting. The GUS staining was performed as described previously (Müller *et al*., 2019), followed by WGA-Alexafluor488 staining to visualize *R. irregularis* (see below). Stained roots were imaged using a BZ-X800LE microscope (Keyence).

### AM fungus staining, microscopy, and quantification of AM fungal colonization

WGA-Alexafluor488 staining was performed by incubating the roots in 20% KOH for 3-4 days at 65°C. Roots were rinsed 3-5 times with distilled water and incubated in 1xPBS overnight. Cleared and washed roots were stained with 0.2 mg/ml WGA-Alexafluor488 (Invitrogen) in 1xPBS for at least 6 hours. Stained roots were visualized using a Leica Thunder Imager or a Leica M205 FA stereomicroscope. Root length colonization was assessed using the grid-line method (McGonigle *et al*., 1990). Arbuscule and vesicle density was quantified as previously described (Müller *et al*., 2020; Chaulagain *et al*., 2023a). In brief, a minimum of 5 z-stacks each of randomly selected colonized root sections from at least 3 independent root systems were imaged that encompassed all arbuscules in that area. Roots were positioned so that the hyphopodium was at the center of the image to ensure infections were at a comparable developmental stage. Imaged arbuscules were counted using the Fiji Image Analysis Package (Schindelin *et al*., 2012), which was also used to accurately measure the imaged root length. Arbuscule density (arbuscules/mm) was calculated by dividing the number of arbuscules by the root length. Confocal images were obtained using a Leica SP5 confocal microscope.

### RNA extraction, cDNA synthesis, and qPCR

RNA extraction, cDNA synthesis, and quantitative PCR were performed as previously described (Müller *et al*., 2019). The qPCRs were performed on a BioRad CFX384 Real-time system. *MtCLE53* gene expression was normalized to the housekeeping gene *MtEF1a* (elongation factor). See Supplementary table 9 for primer sequences.

### Transcriptomics

For each genotype (*crn* and R108), 6-week-old mock-(n=4 per genotype) and *R. irregularis*-colonized root systems (n=3 per genotype) were harvested as described above. WGA-Alexafluor488 staining was performed on a subset of the samples to confirm colonization. RNA was extracted from the remaining samples as previously described (Javot *et al*., 2011). Library preparation and transcriptome sequencing was performed by Novogene Co, Ltd. Samples were enriched for mRNA using poly(A) capture. Paired-end sequencing (PE150) was performed on the Illumina Novaseq X platform. Sequencing raw data are deposited in the NCBI Sequence Read Archive (SRA) (BioProject PRJNA1140176). Filtered reads were mapped against the *M. truncatula* Mt4.0v1 genome (downloaded from Phytozome 13) and *R. irregularis* DAOM RhiirDAOM197198 v2.0 (downloaded from the JGI Mycocosm database) genome using HiSAT2 (Tang *et al*., 2014; Kim *et al*., 2019; Yildirir *et al*., 2022). Normalized read counts were used to calculate differential gene expression with DESeq2 (Love *et al*., 2014). For each pairwise comparison, log2-transformed fold changes were obtained for each mapped gene. Differential gene expression was considered significant if the p-value adjusted for multiple comparisons using Benjamini-Hochberg method (padj) was below 0.05. Heatmaps were constructed using FPKM-normalized read counts; rows were clustered using Pearson correlation, columns using Spearman correlation. To identify gene groups with similar expression patterns across samples, hierarchical clustering was performed on FPKM-normalized read counts averaged for each genotype/treatment. The number of clusters (k=6) was determined using the sum of squared error method. GO term analysis for biological process was performed on https://geneontology.org/; visualization of the obtained GO term enrichment results was performed in R.

### Scanning electron microscopy

For inflorescence phenotyping, plants were grown in 6-inch azalea pots in soil (Promix) under a 16 hour light/8 hour dark regime at 20-22°C and watered with distilled water as needed. Axils from 8-week-old plants were dissected with 30 gauge insulin syringe needles to reveal inflorescence meristems under a stereoscope. Meristems and the surrounding tissue were mounted on stubs using double-sided tape and imaged in a Jeol JCM-7000 benchtop scanning electron microscope (SEM) with the following conditions: high vacuum, 15 kV, high probe current, analysis mode. Meristems from SEM images were categorized based on (Benlloch *et al*., 2003) and measured using Fiji.

## Statistical analyses

Statistical significance between two experimental groups was assessed using a two-sided Student’s t-test. In graphs, statistical significances between two groups are represented as follows: ***p<0.001; **p<0.01; *p<0.05. Statistical significance between multiple experimental groups was calculated with ANOVA, followed by Tukey’s HSD test for multiple comparisons. Pairwise comparisons were considered statistically significant if p<0.05. Significant differences are denoted with different letters. The % reduction of root length colonization shown in Fig. 3 was calculated separately for each genotype by the following formula: ((sample RLC_35S:CLE53_ – average RLC_35S:GUS_)/average RLC_35S:GUS_)*100. The % reduction values for each genotype were then compared to each other using student’s t-test. See paragraphs ‘transcriptomics’ for additional details on the statistics used to analyze RNA sequencing data. Graphs show data from individual experiments. All experiments were repeated at least twice with similar results.

## Acknowledgements

We thank Julia Frugoli (Clemson University, SC) for kindly gifting us the *crn* and sunn-5 mutant lines.

## Author contributions

L.M.M. conceived the project. J.O., E.X.L, M.K., Y.C.T.A., S.B., H.E., C.K.G. and L.M.M. performed all experiments except for the inflorescence meristem analyses, which were performed by P.L.L. L.M.M. and D.J. acquired funding for the project. L.M.M. wrote the manuscript with input from all co-authors.

## Data availability

Raw data from the transcriptome sequencing has been deposited on the NCBI SRA database (Bioproject ID: PRJNA1140176).

