## Supplementary Figures 1-9 for "*Medicago truncatula CORYNE* modulates inflorescence meristem branching, nutrient signaling, and arbuscular mycorrhizal symbiosis"

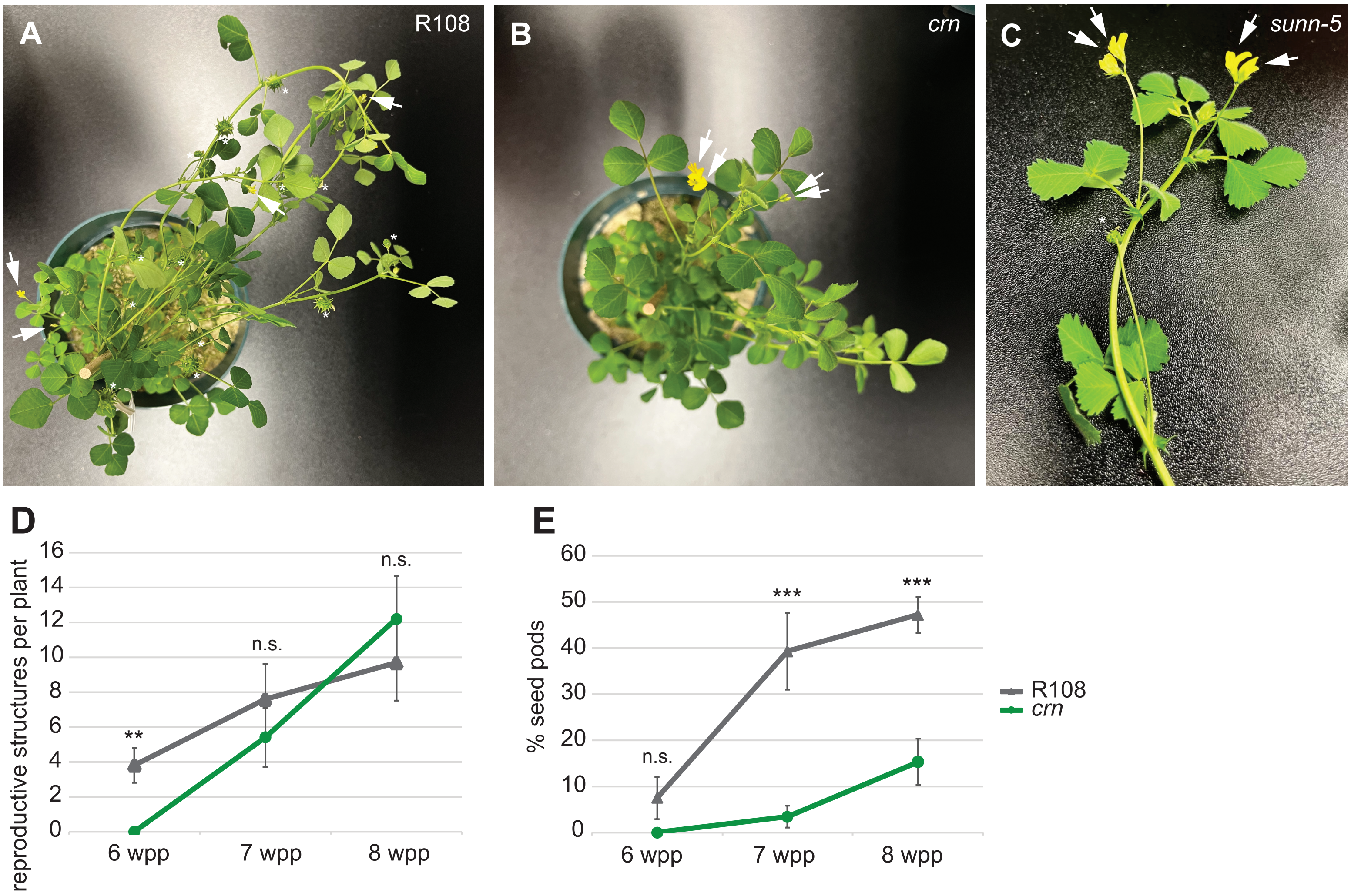


**Supplementary Figure 1: *M. truncatula CRN* regulates flower development.** A) 7 weeks-old R108 wild-type plant with flowers/buds (arrows) and developing or mature seeds (asterisks). Each inflorescence meristem produced a single flower. B) 7 weeks-old *crn* plant with flowers/buds (arrows). The majority of inflorescences produced two flowers. Note, although this plant is the same age as the wild-type plant shown in A), it is generally smaller and no seeds have been produced due to the delay in flowering. C) Fasciated inflorescences (arrows) were also observed in *sunn-5* mutants. D) Reproductive development is delayed in *crn* relative to R108 wild-types. Graph shows the average number of reproductive structures (buds, flowers, pods) per plant over time (6, 7, 8 weeks post planting, wpp). At 6wpp, R108 started to produce reproductive structures, while in *crn* reproductive structures were first observed at 7 wpp. E) Seed pod production is delayed in *crn* plants relative to R108 wild-types. Graph shows the proportion of seed pods out of the total number of reproductive structures shown in panel D). (D, E) Graphs display mean values from 10 individual plants per genotype. Error bars represent standard error. Statistics to compare *crn* and wild-type: t-test, ***$p<0.001$; **$p<0.01$; *$p<0.05$; n.s., not significant.

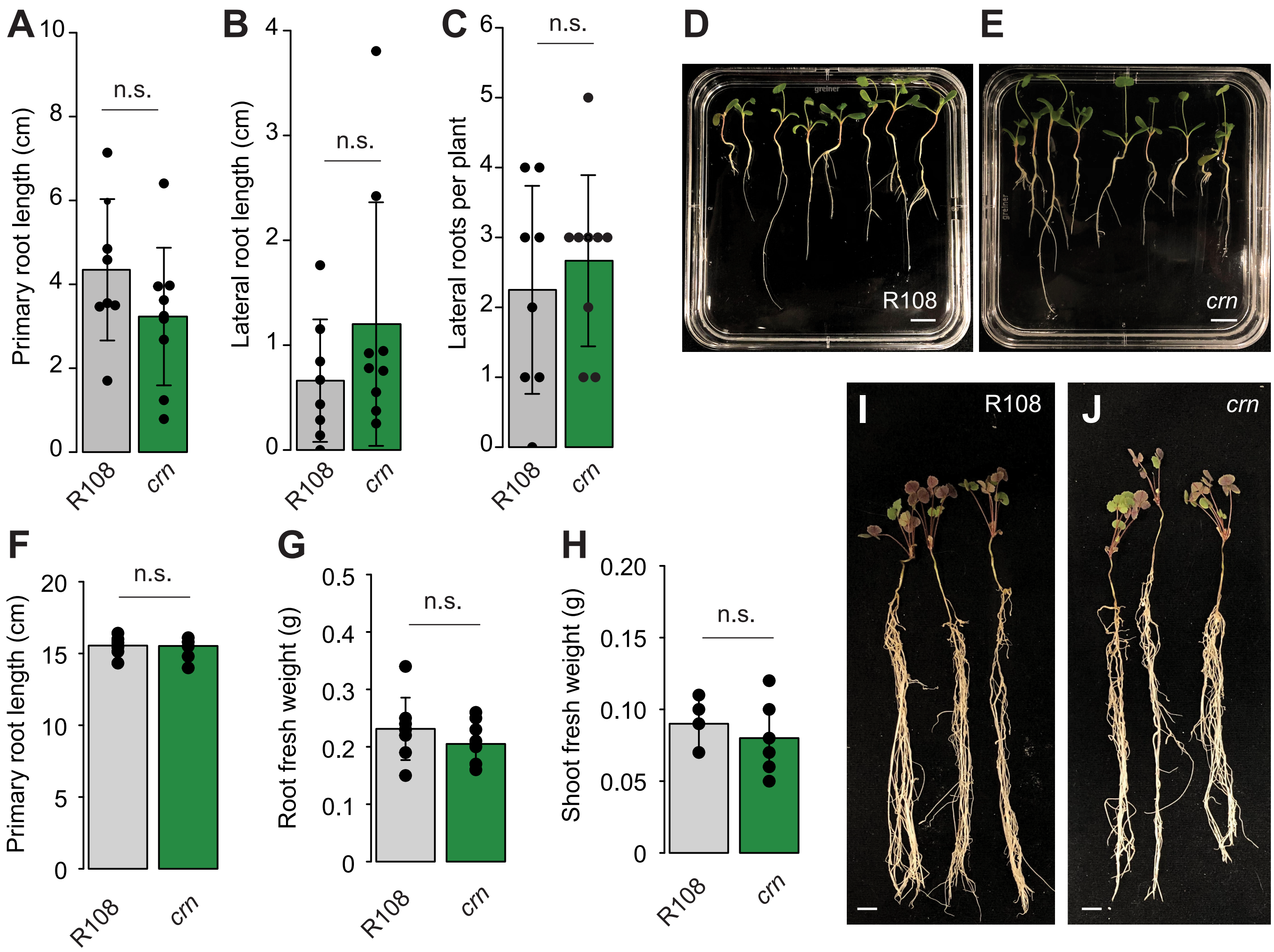


**Supplementary Figure 2: *M*utation in *crn* does not affect root system development.** A) Primary root length of 12-day old *crn* and R108 seedlings grown on F media with 20µM phosphate. B) Lateral root length of 12-day old *crn* and R108 seedlings. C) Number of lateral roots per 12-day old seedling. D, E) Representative images of 12-day old seedlings grown on F media agar plates with 20µM phosphate. F) Primary root length of 4-weeks old *M. truncatula* plants grown in in sand-vermiculite mix under low phosphate conditions. G) Root fresh weight of 4-weeks old plants. H) Shoot fresh weight of 4-weeks old plants. I, J) Representative images of 4-weeks old R108 and *crn* plants grown in sand-vermiculite under a low phosphate fertilization regime in the absence of an AM fungus. Statistical analysis: n.s., not significant (t-test). Scale bars: 1 cm.

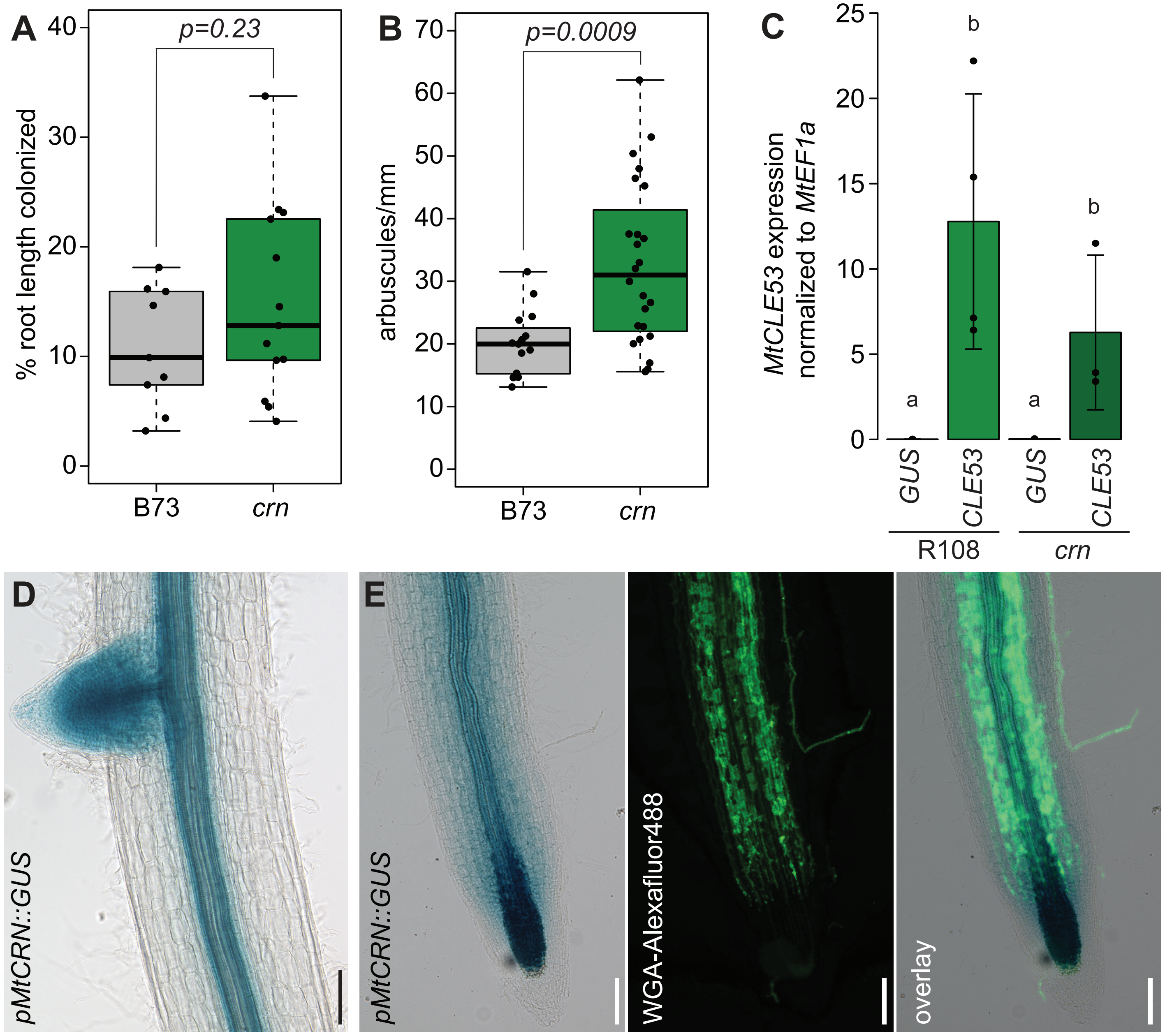


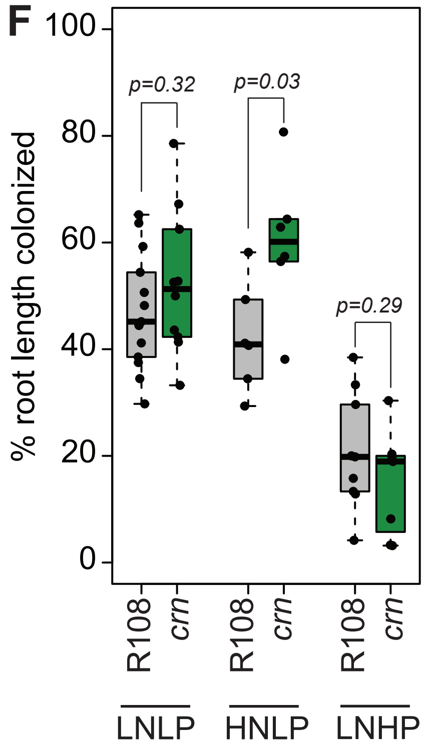


**Supplementary Figure 3: Maize *crn* mutant phenotypes and *pMtCRN::GUS* activity in *M. truncatula* roots.** A) Maize crn mutants show a trend towards increased root colonization by *R. irregularis.* B) Maize *crn* mutants have a higher arbuscule density (arbuscules/mm colonized root) than B73 wildtype controls. Pairwise comparisons in A) and B) were performed using student's t-test; the resulting p-values are shown in each panel. C) *MtCLE53* gene expression in M. truncatula R108 and crn roots transformed with the control construct *35S::GUS* (*GUS*) or *35S::MtCLE53* (*CLE53*). *MtCLE53* overexpression was successful in all samples. Statistics: Anova (p=0.009) followed by Tukey's HSD posthoc test. Different letters denote significant differences (p<0.05) in pairwise comparisons. Colonization data for these samples is shown in Fig. 3. D) Spatial expression pattern of *pMtCRN::GUS* in mock-inoculated *M. truncatula* roots. GUS activity is observed in the emerging lateral roots the vascular tissue. E) Spatial expression pattern of *pMtCRN::GUS* in *R. irregularis*-colonized roots (left). GUS activity is observed in the vascular tissue, the meristemal region of the root, and in and around colonized cortex cells. Middle panel depicts *R. irregularis* counterstaining of the same root with WGA-Alexafluor488. Scale bars in D) and E):100 µm. F) Nutrient manipulation affects *R. irregularis* root length colonization in *M. truncatula crn* mutants. Increased root length colonization is only apparent in high nitrogen/low phosphate (HNLP) conditions, whereas the phenotype is absent in low nitrogen/low phosphate (LNLP) and low nitrogen/high phosphate (LNHP). Student t-tests performed separately for each nutrient treatment (p-values shown in figure). Data compiled from two independent experiments.

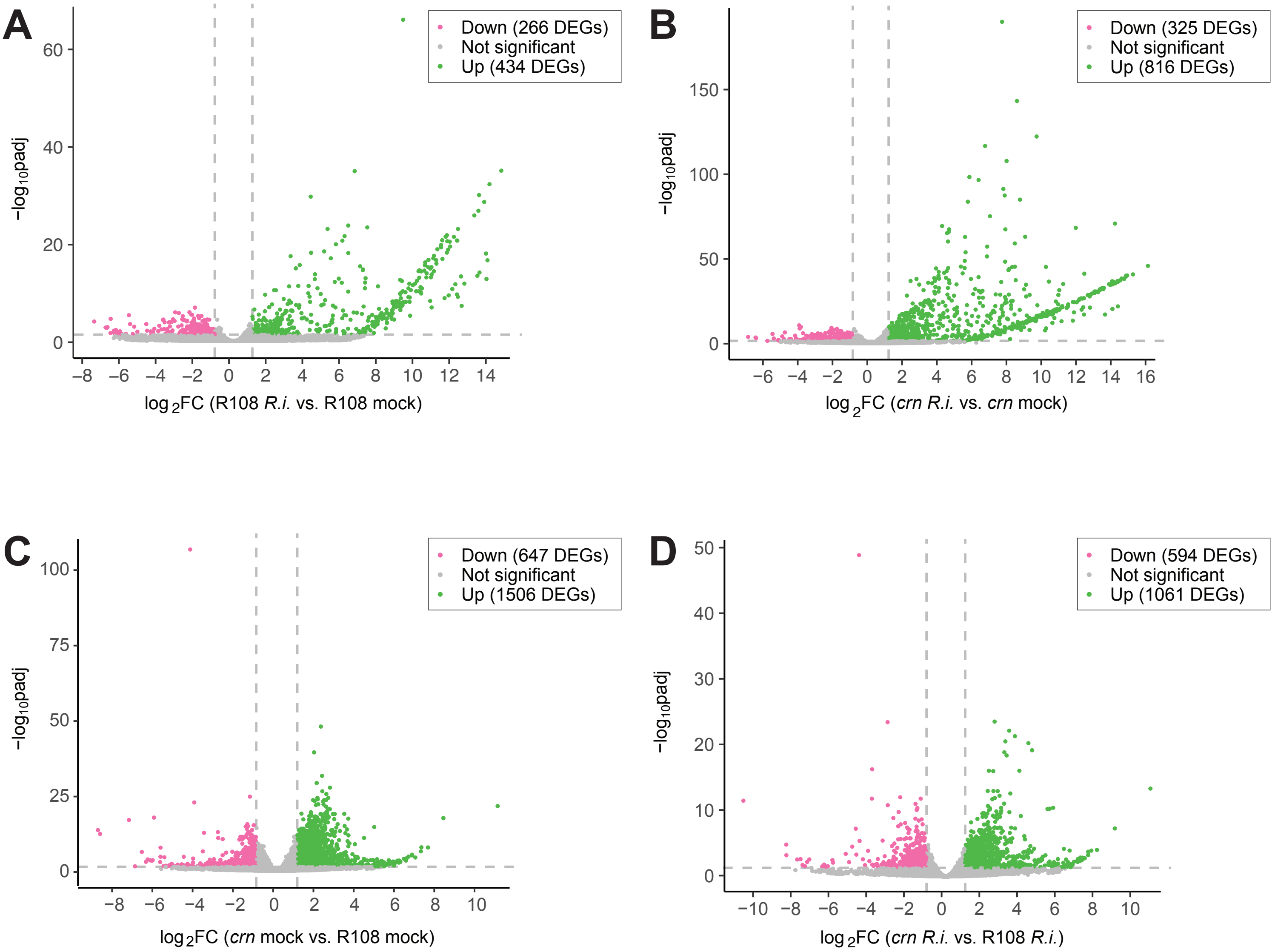


**Supplementary Figure 4: Volcano plots showing global gene expression differences between *crn* and R108 in root transcriptomics data.** (A) *R.irregularis (R.i.)*-inoculated R108 wild-type roots relative to mock-inoculated wild-type roots, (B) *R.irregularis (R.i.)*-inoculated *crn* roots relative to mock-inoculated *crn* roots, (C) mock-inoculated *crn* roots relative to mock-inoculated R108 wild-type roots, and (D) *R.irregularis (R.i.)*-inoculated *crn* roots relative to *R.irregularis (R.i.)*-inoculated R108 wild-type roots. Significantly up-regulated regulated genes (log2FC>1, padj<0.05) in each comparison are shown in green, significantly down-regulated genes (log2FC<-1, padj<0.05) are shown in pink. The number up- or down-regulated DEGs falling within these categories is indicated in each panel. Grey: genes whose exptression is not significantly different in the respecitve comparisons (-1<log2FC<1 or padj>0.05).

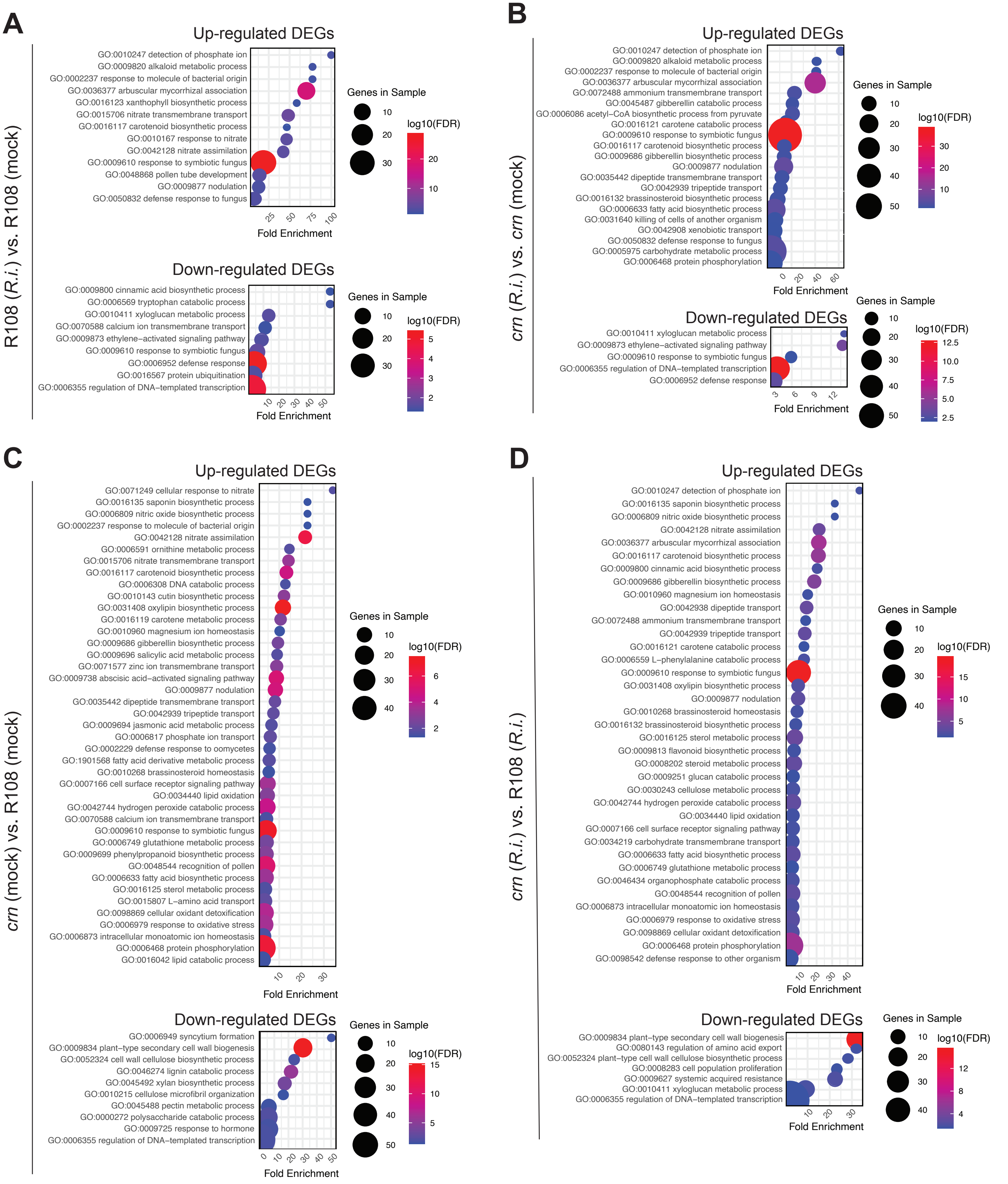


**Supplementary Figure 5: Differentially expressed genes in *crn* and R108 roots**. A) GO terms for 'biological process' of all differentially expressed genes (DEGs; log2 fold-change >1 for up-regulated genes, or <-1 for down-regulated genes; padj<0.05) in R108 roots colonized by *R. irregularis (R.i.)* relative to mock-inoculated roots. B) GO terms corresponding to DEGs in *crn* roots colonized by *R. irregularis (R.i.)* relative to mock-inoculated roots. C) GO terms corresponding to DEGs in mock-inoculated *crn* roots relative to mock-inoculated R108 wild-type roots. D) GO terms corresponding to DEGs in *R. irregularis (R.i.)*-colonized crn roots relative to colonized R108 roots. A-D) The size of the circles corresponds to the number of genes per comparison for each GO term, the color code to the statistical significance (log10 false discovery rate, FDR) of the enrichment.

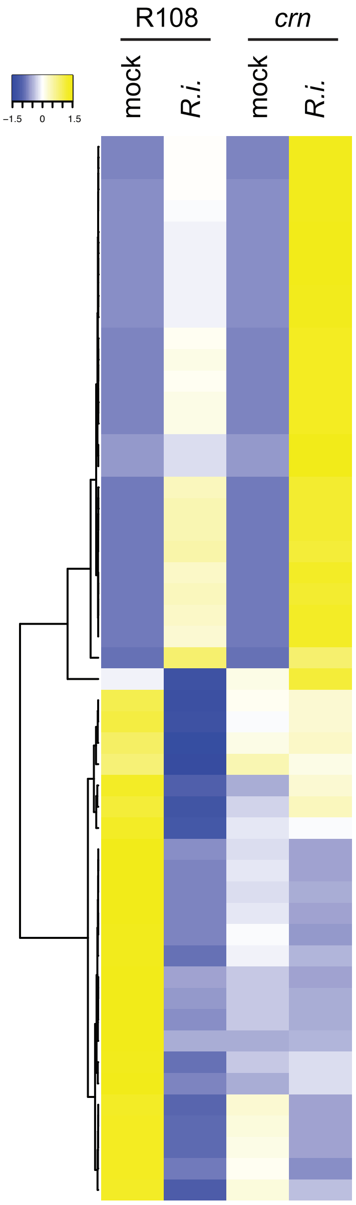


| Gene | log$_2$ fold change: R108 *R.i.* vs. R108 mock | *crn R.i.* vs. *crn* mock | *crn* mock vs. R108 mock | *crn R.i.* vs. R108 *R.i.* |
|---|---|---|---|---|
| *Medtr2g086620 Cupin domain protein* | 11.43*** | 12.86*** | NA | 1.46 |
| *Medtr2g086640 Cupin domain protein* | 11.91*** | 13.34*** | NA | 1.45 |
| *Medtr5g031030 MtLEC5 (lectin)* | 11.87*** | 13.34*** | NA | 1.50* |
| *Medtr8g055940 Chitinase* | 11.86*** | 13.48*** | NA | 1.64 |
| *Medtr3g079570 Serine carboxypeptidase* | 12.08*** | 13.90*** | NA | 1.84* |
| *Medtr1g028600 MtPT4* | 14.22*** | 13.62*** | NA | 1.86*** |
| *Medtr4g118000 Scopoletin glucosyltransferase* | 11.70*** | 13.60*** | NA | 1.92*** |
| *Medtr1g036410 MtAMT2;5* | 11.88*** | 13.63*** | NA | 1.77* |
| *Medtr1g040500 MtRAM2* | 11.92*** | 10.24*** | NA | 1.79*** |
| *Medtr0211s0080 MtKunitz3* | 11.50*** | 12.77*** | NA | 1.29 |
| *Medtr4g077180 MtnsLTP104* | 13.01*** | 14.24*** | NA | 1.26 |
| *Medtr7g068600 MtMYB1* | 13.59*** | 13.46*** | NA | 1.33** |
| *Medtr8g461050 MtKunitz24* | 11.62*** | 12.78*** | NA | 1.19 |
| *Medtr5g031100 Legume lectin domain protein* | 12.26*** | 13.41*** | NA | 1.18 |
| *Medtr2g086630 Germin-like protein* | 11.48*** | 13.77*** | NA | 2.31*** |
| *Medtr0100s0150 MtKunitz2* | 13.04*** | 15.45*** | NA | 2.43*** |
| *Medtr4g076150 MtnsLTP103* | 12.79*** | 13.46*** | NA | 0.70 |
| *Medtr5g035640* | 13.31*** | 13.99*** | NA | 0.70 |
| *Medtr0302s0030 Kunitz trypsin inhibitor* | 13.49*** | 14.12*** | NA | 0.66 |
| *Medtr8g059790 MtKPI106\|MtKunitz22* | 13.06*** | 13.58*** | NA | 0.54 |
| *Medtr3g014820 MtKunitz7* | 12.95*** | 13.77*** | NA | 0.84 |
| *Medtr8g012805 MtDefMd1* | 13.44*** | 14.21*** | NA | 0.80 |
| *Medtr5g075520* | 13.41*** | 14.29*** | NA | 0.91 |
| *Medtr7g086220 MtPCY19\|MtSC18* | 11.78*** | 12.76*** | NA | 1.01 |
| *Medtr4g091020 LysM-domain protein* | 12.12*** | 12.10*** | NA | 0.01 |
| *Medtr3g035600 Photosystem I psaA/psaB protein* | -6.35** | 0.75 | 0.27 | 7.35*** |
| *Medtr4g088955 Asparagine synthetase* | -6.18* | 0.16 | -0.75 | 5.59 |
| *Medtr1g101020 NOTUM-RELATED* | -6.31* | 0.27 | -0.91 | 5.67 |
| *Medtr2g091120* | -6.32* | 0.16 | -0.58 | 5.90 |
| *Medtr2g035430 Cationic amino acid transporter* | -5.98* | -0.27 | -0.22 | 5.47 |
| *Medtr7g078640 Leucine-rich repeat containing protein* | -5.72* | 1.27 | -2.08 | 4.90 |
| *Medtr2g021540* | -6.10* | 0.77 | -1.35 | 5.51 |
| *Medtr5g049780 PPR repeat family protein* | -5.96* | 0.12 | -1.22 | NA |
| *Medtr7g105170 No apical meristem (NAM)* | -6.52*** | -1.67 | -2.11 | 2.77 |
| *Medtr7g023740 Polygalacturonase inhibitor* | -7.40*** | -1.73 | -1.97 | NA |
| *Medtr5g080000 Leucine-rich repeat containing protein* | -6.37* | -1.00 | -2.13 | NA |
| *Medtr5g081360* | -5.28** | -1.60 | -1.73 | NA |
| *Medtr5g082270 Protein kinase* | -6.69** | -2.22 | -1.62 | NA |
| *Medtr1g116100* | -6.79** | -0.75 | -1.59 | NA |
| *Medtr8g432450 BON1-ASSOCIATED PROTEIN* | -5.56*** | -3.08 | -2.92* | NA |
| *Medtr3g465090 S-linalool synthase* | -6.53* | NA | -3.02 | NA |
| *Medtr8g013780 Cation transporting ATPase* | -5.49*** | -0.74 | -2.42** | 2.34 |
| *Medtr1g115890 Chalcone isomerase* | -6.25** | NA | -5.67*** | NA |
| *Medtr0032s0070* | -5.35* | 0.17 | -1.87 | NA |
| *Medtr7g076620* | -6.09* | 1.09 | -3.09* | NA |
| *Medtr6g037010 Protein of unknown function (DUF793)* | -5.93* | -1.58 | -0.87 | NA |
| *Medtr5g014310 Aluminium-activated malate transporter* | -6.18* | -1.79 | -0.99 | NA |
| *Medtr7g069580 NB-ARC domain protein* | -6.21** | -1.39 | -1.09 | NA |
| *Medtr1g110870 MtCONSTANS-like* | -6.03* | -2.37 | -1.19 | NA |
| *Medtr4g091840 G-type lectin S-RLK* | -6.72** | -0.97 | -0.83 | NA |

**Supplementary Figure 6: Top 25 DEGs in *R. irregularis (R.i.)*-colonized R108 relative to mock-inoculated R108 roots.** Heatmap shows averages of n=3 samples for *R. irregularis*-inoculated R108 and *crn* roots and n=4 samples for mock-inoculated R108 and *crn* roots, respectively, including log2-transformed fold changes for each comparison. Asterisks indicate statistical significance for each comparison: ***padj<0.05; **padj<0.01; *padj<0.05. Abbreviations: *MtPT4, PHOSPHATE TRANSPORTER 4; MtDefMd1, ENOD-like 29|mycorrhiza-dependent defensin 1; MtKPI* or *MtKuniz, Kunitz-type trypsin inhibitor; MtnsLTP104, nonspecific lipid transfer protein; MtPCY19|MtSC18, Plantacyanin/Chemocyanin 19|stellacyanin 18; RLK, receptor-like kinase.*

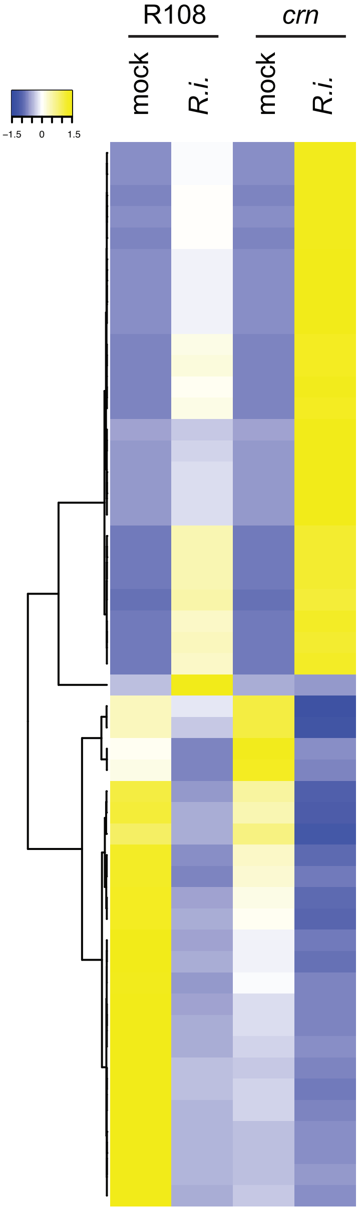


| Gene | log$_2$ fold change R108 *R.i.* vs. R108 mock | *crn R.i.* vs. *crn* mock | *crn* mock vs. R108 mock | *crn R.i.* vs. R108 *R.i.* |
|---|---|---|---|---|
| *Medtr5g031090 Legume lectin protein* | 10.44*** | 13.45*** | NA | 1.58 |
| *Medtr8g055940 Chitinase* | 11.86*** | 13.48*** | NA | 1.64 |
| *Medtr6g034940 MtCYP733A1* | 8.08** | 13.00*** | NA | 1.48 |
| *Medtr5g031030 MtLEC5* | 11.87*** | 13.34*** | NA | 1.50* |
| *Medtr2g086640 Cupin domain protein* | 11.91*** | 13.34*** | NA | 1.45 |
| *Medtr1g028600 MtPT4* | 14.22*** | 13.62*** | NA | 1.86*** |
| *Medtr3g079570 Serine carboxypeptidase* | 12.08*** | 13.90*** | NA | 1.84* |
| *Medtr4g118000 Scopoletin glucosyltransferase* | 11.70*** | 13.60*** | NA | 1.92*** |
| *Medtr1g036410 MtAMT2;5* | 11.88*** | 13.63*** | NA | 1.77* |
| *Medtr5g031100 Legume lectin protein* | 12.26*** | 13.41*** | NA | 1.18 |
| *Medtr8g068030 MtLEC7* | NA | 14.62*** | NA | 1.09 |
| *Medtr7g068600 MtMYB1* | 13.59*** | 13.46*** | NA | 1.33** |
| *Medtr4g077180 MtnsLTP104* | 13.01*** | 14.24*** | NA | 1.26 |
| *Medtr4g047610 MtCP3/AtSAG12-like* | 10.01*** | 13.06*** | NA | 3.08* |
| *Medtr5g022560 MtCP4/MtCP2* | 11.09*** | 13.78*** | NA | 2.71*** |
| *Medtr3g079190 Neutral ceramidase* | 11.08*** | 13.28*** | NA | 2.22*** |
| *Medtr2g086630 Germin-like protein* | 11.48*** | 13.77*** | NA | 2.31*** |
| *Medtr0100s0150 MtKunitz2* | 13.04*** | 15.45*** | NA | 2.43*** |
| *Medtr4g076150 MtnsLTP103* | 12.79*** | 13.46*** | NA | 0.70 |
| *Medtr5g035640* | 13.31*** | 13.99*** | NA | 0.70 |
| *Medtr0302s0030 Kunitz trypsin inhibitor* | 13.49*** | 14.12*** | NA | 0.66 |
| *Medtr8g059790 MtKPI106\|MtKunitz22* | 13.06*** | 13.58*** | NA | 0.54 |
| *Medtr3g014820 MtKunitz7* | 12.95*** | 13.77*** | NA | 0.84 |
| *Medtr8g012805 MtDefMd1* | 13.44*** | 14.21*** | NA | 0.80 |
| *Medtr5g075520* | 13.41*** | 14.29*** | NA | 0.91 |
| *Medtr4g024265 Ribosomal protein S15P/S13E* | NA | -5.14* | -0.81 | NA |
| *Medtr5g058470 AP2/ERF transcription factor* | -0.48 | -4.17** | NA | -3.24 |
| *Medtr4g059520* | -0.86 | -5.76* | 0.52 | NA |
| *Medtr4g017720 Disease resistance protein* | -4.79 | -5.15* | 1.31 | NA |
| *Medtr4g036880* | NA | -6.42*** | 1.19 | NA |
| *Medtr2g089210 EF-hand domain protein* | -2.19* | -3.90* | -0.39 | -2.10 |
| *Medtr8g432370 BON1-ASSOCIATED PROTEIN* | -1.74 | -3.96*** | -0.53 | -2.76** |
| *Medtr5g464340* | -1.78 | -6.83*** | -0.17 | NA |
| *Medtr8g432350 BON1-ASSOCIATED PROTEIN* | -2.68** | -3.98*** | -0.75 | -2.05 |
| *Medtr8g079950 Transferase* | -3.98 | -4.10* | -0.91 | NA |
| *Medtr4g128410 Xyloglucan endotransglucosylase* | -2.36 | -3.96* | -0.98 | NA |
| *Medtr3g116720 DNJ-23-related* | -2.21 | -5.09* | -1.10 | NA |
| *Medtr2g043050 AP2/ERF transcription factor* | -3.00 | -4.16** | -1.61 | -2.78* |
| *Medtr4g128570 Xyloglucan endotransglucosylase* | -2.60*** | -4.78*** | -1.61 | -3.81*** |
| *Medtr0008s0470 MtMYB014* | -3.72*** | -3.89*** | -1.59 | -1.75 |
| *Medtr2g043030 AP2/ERF transcription factor* | -3.24** | -4.21*** | -1.92* | -2.89 |
| *Medtr3g434920 Copper transporter* | -3.38** | -6.39** | -2.07* | NA |
| *Medtr4g128590 Xyloglucan endotransglucosylase* | -3.62* | -4.19* | -2.28 | -2.86** |
| *Medtr4g086165 AP2/ERF transcription factor* | -2.48** | -4.84** | -2.28 | -4.65*** |
| *Medtr1173s0010 Copper transporter* | -2.40 | -5.09* | -2.18 | NA |
| *Medtr2g043040 AP2/ERF transcription factor* | -2.81 | -5.40** | -2.28 | -4.88 |
| *Medtr2g097390 GRAS domain family protein* | -2.98 | -4.44** | -2.81* | -4.29** |
| *Medtr2g097310 GRAS domain family protein* | -3.10*** | -4.93*** | -2.78* | -4.62*** |
| *Medtr4g086160 AP2/ERF transcription factor* | -3.34*** | -5.47*** | -3.03* | -5.17*** |
| *Medtr4g086190 AP2/ERF transcription factor* | -3.45 | -4.53** | -2.68 | -3.77* |

**Supplementary Figure 7: Top 25 DEGs in *R. irregularis (R.i.)*-colonized *crn* roots relative to mock-inoculated *crn* roots.** Heatmap shows averages of n=3 samples for *R. irregularis*-inoculated R108 and *crn* roots and n=4 samples for mock-inoculated R108 and *crn* roots, respectively, including log2-transformed fold changes for each comparison. Asterisks indicate statistical significance for each comparison: ***padj<0.05; **padj<0.01; *padj<0.05. Abbreviations: *MtPT4, PHOSPHATE TRANSPORTER 4; MtAMT2;5, AMMONIUM TRANSPORTER 2;5; MtCP4/MtCP2, Cysteine protease; MtCP3/AtSAG12-like, cystein protease 3/senescence-associated gene 12, MtKPI* or *MtKuniz, Kunitz-type trypsin inhibitor; MtDefMd1, ENOD-like 29*\|*mycorrhiza-dependent defensin 1.*

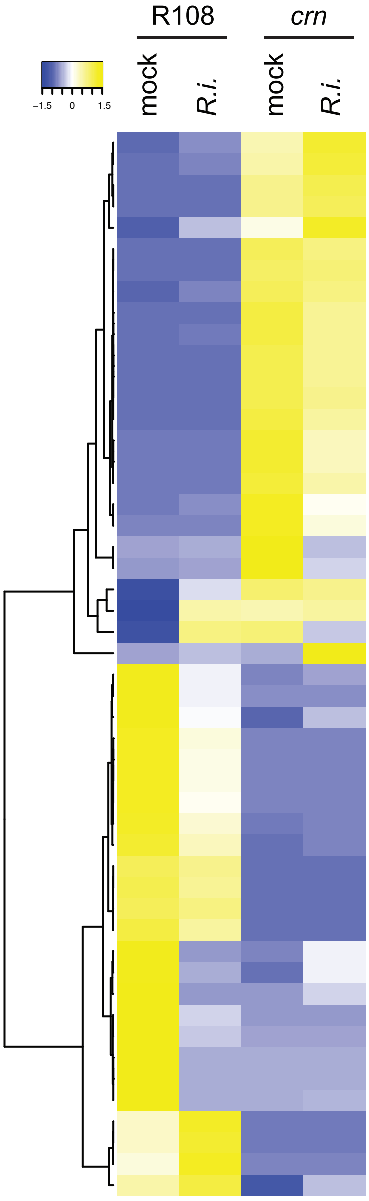


| | log$_2$ fold change | | | |
|---|---|---|---|---|
| Gene | R108 *R.i.* vs. R108 mock | *crn R.i.* vs. *crn* mock | *crn* mock vs. R108 mock | *crn R.i.* vs. R108 *R.i.* |
| *Medtr7g056670* | NA | 0.66 | 5.84** | 3.34 |
| *Medtr1g006605* | NA | 0.54 | 5.88*** | 3.97 |
| *Medtr8g089342 Amino acid permease* | NA | 0.32 | 6.30*** | 6.55* |
| *Medtr7g104390 Phosphatidylserine decarboxylase* | NA | 0.24 | 6.35*** | 6.53* |
| *Medtr8g074170 FI18644P1-RELATED* | NA | 0.91 | 5.78** | 1.78 |
| *Medtr3g058320* | NA | -0.20 | 6.42*** | 6.16* |
| *Medtr3g067615 F-box protein* | NA | -0.14 | 10.77*** | 10.57*** |
| *Medtr2g426020 F-box protein* | NA | -0.23 | 6.20*** | 2.93 |
| *Medtr7g090270 Unknown function DUF1589* | NA | -0.42 | 6.59*** | 6.11* |
| *Medtr3g011930 Serine/threonine RLK* | NA | -0.41 | 8.12*** | 5.37*** |
| *Medtr4g108720 AGAMOUS-like* | NA | -0.39 | 5.82** | 5.37 |
| *Medtr2g101310 Ankyrin repeat-containing protein* | NA | -0.39 | 7.37*** | 6.92** |
| *Medtr3g101450 DNA-directed RNA polymerase beta* | NA | -0.34 | 7.03*** | 6.63** |
| *Medtr8g042355* | NA | -0.48 | 6.76*** | 6.22* |
| *Medtr4g013245 MtFPN3* | NA | -0.76 | 5.97*** | NA |
| *Medtr5g056735 WRKY transcription factor* | NA | -2.67 | 5.95** | NA |
| *Medtr7g104380 Glucose-1-P adenylyltransferase* | NA | -0.62 | 6.60*** | 5.92 |
| *Medtr7g098630 Subtilisin-like protease* | NA | -1.23 | 5.70** | NA |
| *Medtr4g114080 Lactoylglutathione lyase glyoxalase* | NA | -1.12 | 6.02*** | NA |
| *Medtr2g438320* | NA | -3.23 | 5.71* | NA |
| *Medtr2g075750 WRKY transcription factor* | NA | -2.67 | 5.95** | NA |
| *Medtr4g104870* | NA | -0.13 | 5.82** | 0.78 |
| *Medtr2g044090 cAMP response element binding* | 5.91 | 0.07 | 5.87** | 0.04 |
| *Medtr0552s0040 LRR domain containing protein* | 5.72 | -1.20 | 5.77** | -1.16 |
| *Medtr2g101920 Lysine histidine transporter* | 8.84*** | 5.37*** | 7.05*** | 3.57*** |
| *Medtr5g040970* | -1.68 | NA | -6.08*** | NA |
| *Medtr5g008790 AP2/ERF transcription factor* | -1.79 | NA | -6.29*** | NA |
| *Medtr3g082830 Zinc finger FYVE domain* | -1.31 | NA | -5.26* | NA |
| *Medtr3g050710 Pectinesterase* | -1.16 | NA | -5.69*** | NA |
| *Medtr5g086750 Myb/SANTlike DNAbinding domain* | -1.19 | NA | -8.72*** | -8.24*** |
| *Medtr6g466310* | -1.16 | NA | -5.67*** | -3.80* |
| *Medtr2g038680* | -1.28 | 0.53 | -5.99*** | -4.18* |
| *Medtr8g055900* | -0.92 | NA | -6.58*** | -4.50 |
| *Medtr1g080080 Myb/SANT-like DNA-binding* | -0.67 | NA | -5.08* | -2.66 |
| *Medtr1g102335 Mucin-related protein* | -0.28 | NA | -6.05*** | -5.76 |
| *Medtr0430s0010 Lactoylglutathione lyase* | -0.35 | NA | -8.61*** | -8.24*** |
| *Medtr4g045750* | -0.16 | NA | -6.28*** | -6.10* |
| *Medtr1g046650 E3 ubiquitin ligase* | -0.48 | NA | -5.41* | NA |
| *Medtr4g037660* | -4.14 | NA | -5.14* | NA |
| *Medtr7g092980* | -2.73 | NA | -5.35* | NA |
| *Medtr2g097650* | -5.18 | NA | -5.22* | NA |
| *Medtr3g108960 WRKY transcription factor* | -2.40 | NA | -5.48*** | -3.13 |
| *Medtr3g109860 Unknown function DUF1677* | -3.12 | NA | -6.91* | NA |
| *Medtr2g089665* coumaryl transferase | -5.11 | NA | -5.14* | NA |
| *Medtr3g105480 AP2/ERF transcription factor* | -5.48 | NA | -5.52* | NA |
| *Medtr1g115890 Chalcone isomerase* | -6.25** | NA | -5.67*** | NA |
| *Medtr4g036245 Betaglucuronosyltransferase* | 0.92 | NA | -6.16*** | -7.05** |
| *Medtr8g080060* | 0.83 | NA | -7.21*** | -10.46 |
| *Medtr4g117120 Dual specificity protein phosphatase* | 1.15 | NA | -6.38** | -7.51** |
| *Medtr1g084950 AGAMOUS-like MADS-BOX protein* | 0.43 | 4.04 | -5.12** | -1.51 |

**Supplementary Figure 8: Top 25 DEGs in mock-inoculated *crn* relative to mock-inoculated R108 roots.** Heatmap shows averages of n=3 samples for *R. irregularis*-inoculated R108 and *crn* roots and n=4 samples for mock-inoculated R108 and *crn* roots, respectively, including log2-transformed fold changes for each comparison. Asterisks indicate statistical significance for each comparison: ***padj<0.05; **padj<0.01; *padj<0.05. Abbreviations: *MtFPN3, FERROPORTIN 3; coumaryl transferase, COUMAROYL-COA:ANTHOCYANIDIN 3-O-GLUCOSIDE-6''-O-COUMAROYLTRANSFERASE.*

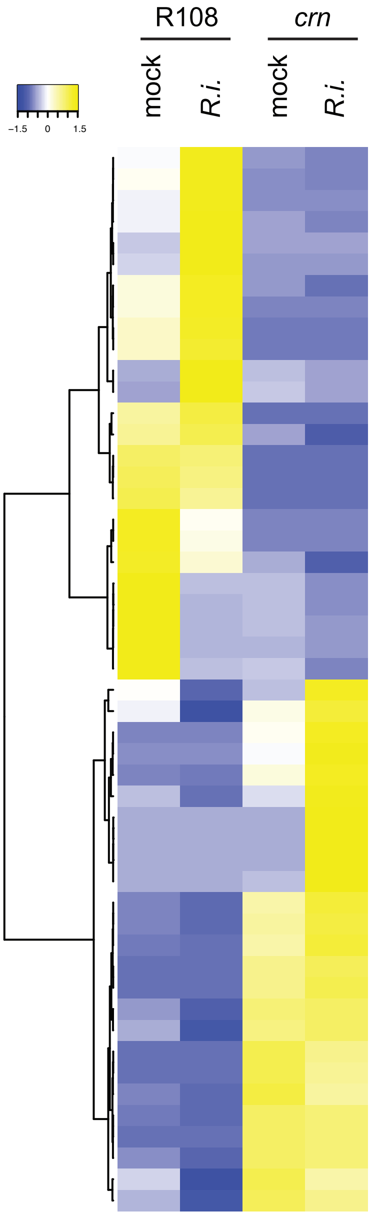


| Gene | log₂ fold change: R108 *R.i.* vs. R108 mock | *crn R.i.* vs. *crn* mock | *crn* mock vs. R108 mock | *crn R.i.* vs. R108 *R.i.* |
|---|---|---|---|---|
| *Medtr0929s0020 Trichome birefringence-like* | 1.55 | NA | -2.70 | -6.19* |
| *Medtr3g014430* | 1.36 | NA | -3.34 | -7.17** |
| *Medtr2g089040 Formin-like protein* | 1.63 | 0.24 | -3.07*** | -4.46*** |
| *Medtr8g070940 MtWIP1a* | -1.76 | NA | 1.63 | -7.28* |
| *Medtr3g093940* | 2.77 | NA | -2.53 | -5.86** |
| *Medtr7g110990 Putative expansin* | NA | NA | -5.11 | -7.68** |
| *Medtr4g094328 MtVTL5* | 1.05 | -2.63 | -2.16* | -5.83** |
| *Medtr4g117120 Dual specificity protein phosphatase* | 1.15 | NA | -6.38** | -7.51** |
| *Medtr4g036245 Betaglucuronosyltransferase* | 0.92 | NA | -6.16*** | -7.05** |
| *Medtr8g080060* | 0.83 | NA | -7.21*** | -10.46*** |
| *Medtr3g085705 Peptidyl-Prolyl cis-trans isomerase C* | 5.22 | NA | NA | -6.29* |
| *Medtr1g103690 Cysteine synthase* | 7.45* | -4.43 | 4.47 | -7.40* |
| *Medtr3g109260 Mago nashi protein* | 0.41 | -0.07 | -3.55*** | -4.02*** |
| *Medtr2g013920 Transcription factor bHLH162* | 0.30 | -4.46 | -1.63 | -6.38* |
| *Medtr7g020850 MtCRN* | -0.07 | -0.33 | -4.22*** | -4.48*** |
| *Medtr4g045750* | -0.16 | NA | -6.28*** | -6.10* |
| *Medtr0430s0010 Lactoylglutathione lyase* | -0.35 | NA | -8.61*** | -8.24*** |
| *Medtr2g038680* | -1.28 | 0.53 | -5.99*** | -4.18* |
| *Medtr5g086750 Myb/SANTlike DNAbinding domain* | -1.19 | NA | -8.72*** | -8.24*** |
| *Medtr4g017700 LRR receptor-like protein* | -0.78 | -3.66* | -1.94* | -4.84*** |
| *Medtr2g097390 GRAS domain family protein* | -2.98 | -4.44** | -2.81* | -4.29** |
| *Medtr2g097310 GRAS domain family protein* | -3.10*** | -4.93*** | -2.78* | -4.62*** |
| *Medtr4g086160 AP2/ERF transcription factor* | -3.34*** | -5.47*** | -3.03* | -5.17*** |
| *Medtr4g086220 AP2/ERF transcription factor* | -3.75*** | -5.00 | -3.72* | -5.00** |
| *Medtr4g086165 AP2/ERF transcription factor* | -2.48** | -4.84** | -2.28 | -4.65*** |
| *Medtr1g026140 MtGST* | -5.61 | 2.03 | -0.81 | 6.82** |
| *Medtr3g035600 Photosystem I psaA/psaB protein* | -6.35** | 0.75 | 0.27 | 7.35*** |
| *Medtr1g075865 Strictosidine synthase* | NA | 1.34 | NA | 7.02** |
| *Medtr1g052170 TMV resistance protein* | NA | 1.61 | 4.81** | 7.81*** |
| *Medtr3g081630* $Fe^{2+}$*transport protein, Zn transporter* | NA | 1.03 | 3.42* | 7.17** |
| *Medtr1g048360 Legume lectin domain* | -5.12 | 1.74 | 0.50 | 7.35*** |
| *Medtr3g079783 Disease resistance protein* | NA | 6.51*** | NA | 6.48* |
| *Medtr3g438050 Polygalacturonase* | NA | 7.09*** | NA | 7.05** |
| *Medtr2g093750 MLO-like protein* | NA | 6.08*** | 0.17 | 8.73*** |
| *Medtr4g107190* | NA | 4.08* | NA | 6.50* |
| *Medtr2g045100 Auxin binding protein* | NA | 0.49 | 2.80 | 6.87** |
| *Medtr5g084070 Fbox/FBD/LRRrepeat protein* | NA | 0.46 | 3.34** | 7.52*** |
| *Medtr5g035240 Disease resistance protein* | NA | 0.55 | 4.89*** | 7.36*** |
| *Medtr7g104390 Phosphatidylserine decarboxylase* | NA | 0.24 | 6.35*** | 6.53* |
| *Medtr8g089342 Amino acid permease* | NA | 0.32 | 6.30*** | 6.55* |
| *Medtr6g083000 LRK10L protein* | -4.80 | 0.07 | 2.09 | 6.95* |
| *Medtr3g020230 G-type lectin S-RLK* | -5.32 | 0.14 | 1.61 | 7.06** |
| *Medtr3g101450 DNA-directed RNA polymerase beta* | NA | -0.34 | 7.03*** | 6.63** |
| *Medtr2g101310 Ankyrin repeat-containing protein* | NA | -0.39 | 7.37*** | 6.92** |
| *Medtr7g098780 Pleiotropic drug resistance protein* | NA | -0.47 | 3.45*** | 6.83** |
| *Medtr5g061290 E3 ubiquitin protein ligase* | NA | -0.09 | 3.82*** | 6.95** |
| *Medtr3g067615 F-box protein* | NA | -0.14 | 10.77*** | 10.57*** |
| *Medtr4g021095 Disease resistance protein* | NA | -0.12 | 2.74 | 6.81** |
| *Medtr3g092320* | -5.95 | -0.38 | 1.23 | 6.79** |
| *Medtr0549s0010 S-type anion channel* | -5.41 | -0.21 | 1.61 | 6.81** |

**Supplementary Figure 9: Top 25 DEGs in *R. irregularis (R.i.)*-inoculated *crn* roots relative to *R. irregularis (R.i.)*-inoculated R108 roots.** Heatmap shows averages of n=3 samples for *R. irregularis*-inoculated R108 and *crn* roots and n=4 samples for mock-inoculated R108 and *crn* roots, respectively, including log2-transformed fold changes for each comparison. Asterisks indicate statistical significance for each comparison: ***padj<0.05; **padj<0.01; *padj<0.05. Abbreviations: *MtGST, Glutathione S-transferase; MtVTL5, Vacuolar iron Transporter-Like protein 5; MtCRN, CORYNE, MtWIP1a, tryptophan-proline-proline domain-interacting protein 1a; MtLRK10L, LEAF RUST 10 DISEASE RESISTANCE LOCUS RECEPTOR-LIKE PROTEIN KINASE-like.*
